# Validation of new bioinformatic tools to identify expanded repeats: a non-reference intronic pentamer expansion in *RFC1* causes CANVAS

**DOI:** 10.1101/597781

**Authors:** Haloom Rafehi, David J Szmulewicz, Mark F Bennett, Nara LM Sobreira, Kate Pope, Katherine R Smith, Greta Gillies, Peter Diakumis, Egor Dolzhenko, Michael A Eberle, María García Barcina, David P Breen, Andrew M Chancellor, Phillip D Cremer, Martin B. Delatycki, Brent L Fogel, Anna Hackett, G. Michael Halmagyi, Solange Kapetanovic, Anthony Lang, Stuart Mossman, Weiyi Mu, Peter Patrikios, Susan L Perlman, Ian Rosemargy, Elsdon Storey, Shaun RD Watson, Michael A Wilson, David Zee, David Valle, David J Amor, Melanie Bahlo, Paul J Lockhart

**Author notes:** Correspondence to Dr Paul Lockhart, Murdoch Children’s Research Institute, Flemington Rd, Parkville, Victoria 3052, Australia. These authors contributed equally to this work (Haloom Rafehi, David J Szmulewicz). These authors contributed equally to this work (Melanie Bahlo, Paul J Lockhart).

## Abstract

Genomic technologies such as Next Generation Sequencing (NGS) are revolutionizing molecular diagnostics and clinical medicine. However, these approaches have proven inefficient at identifying pathogenic repeat expansions. Here, we apply a collection of bioinformatics tools that can be utilized to identify either known or novel expanded repeat sequences in NGS data. We performed genetic studies of a cohort of 35 individuals from 22 families with a clinical diagnosis of cerebellar ataxia with neuropathy and bilateral vestibular areflexia syndrome (CANVAS). Analysis of whole genome sequence (WGS) data with five independent algorithms identified a recessively inherited intronic repeat expansion [(AAGGG)_exp_] in the gene encoding Replication Factor C1 (*RFC1*). This motif, not reported in the reference sequence, localized to an Alu element and replaced the reference (AAAAG)_11_ short tandem repeat. Genetic analyses confirmed the pathogenic expansion in 18 of 22 CANVAS families and identified a core ancestral haplotype, estimated to have arisen in Europe over twenty-five thousand years ago. WGS of the four *RFC1* negative CANVAS families identified plausible variants in three, with genomic re-diagnosis of SCA3, spastic ataxia of the Charlevoix-Saguenay type and SCA45. This study identified the genetic basis of CANVAS and demonstrated that these improved bioinformatics tools increase the diagnostic utility of WGS to determine the genetic basis of a heterogeneous group of clinically overlapping neurogenetic disorders.

## INTRODUCTION

Repetitive DNA sequences constitute approximately one third of the genome and are thought to contribute to diversity within and between species.^1^ Microsatellites or short tandem repeats (STRs) are mini-repeats of DNA, typically two to five base-pairs in length, which are usually present in a concatamer of between five and fifty repeated elements. There are thousands of STRs scattered through the human genome and recent studies have suggested important roles for STRs in the regulation of gene expression.^2;3^ STRs display considerable variability in length between individuals, which is presumed to have no detrimental consequences for humans^4;5^ unless the repeat number is expanded beyond a gene-specific threshold.^6;7^ Pathogenic repeat expansions (REs) have been shown to underlie at least 30 inherited human diseases, the majority being disorders of the nervous system.^8^ These disorders, which variably have autosomal dominant, autosomal recessive and X-linked inheritance, have an overall prevalence of ∼1:20,000.^9^ They display a broad onset age and are characterized by progressive cerebellar ataxia with dysarthria, oculomotor abnormalities, cognitive dysfunction and other symptoms.^10^ Additional novel pathogenic REs likely remain to be identified. For example, putative spinocerebellar ataxia (SCA) loci, including SCA25 (MIM: 608703) and SCA30 (MIM: 613371) remain to be identified, and unsolved hereditary ataxias such as cerebellar ataxia with neuropathy and bilateral vestibular areflexia syndrome (CANVAS, MIM: 614575) display extensive clinical similarities with known RE disorders.

CANVAS is a cerebellar ataxia with combined cerebellar, vestibular and somatosensory dysfunction.^11;12^ Historically, individuals with CANVAS have been assigned the diagnosis of idiopathic late onset cerebellar ataxia.^13^ More recently, CANVAS is clinically recognized and has been incorporated into the contemporary research and teaching of both cerebellar and vestibular diseases.^14;15^ Unifying the oto-and neuropathology, CANVAS is a neuronopathy (ganglionopathy) affecting the vestibular^16^ and dorsal root ganglia.^17^ The progression of these clinical features can be measured longitudinally using a specific neurophysiological protocol.^18^ A characteristic radiological pattern of cerebellar atrophy has also been described and verified on post-mortem pathology.^11^ The characteristic oculomotor abnormality seen in combined cerebellar and vestibular impairment is the visually-enhanced vestibulo-ocular reflex (VVOR), and this can now be evaluated using a commercially available instrumented assessment tool.^19-21^ Altogether, these advances have allowed the formulation of diagnostic criteria to aid identification of CANVAS, contributing both research and clinical benefits including improved prognostication and targeted management.^12;14^ While detailed clinical findings have driven gene discovery in RE disorders such as Friedreich taxia^22^ the underlying genetic cause(s) of CANVAS has, until very recently, remained elusive (see below).

The majority of individuals and families with CANVAS have been identified in individuals of European ancestry, although CANVAS has recently been reported in two individuals of Japanese ethnicity, a 68-year old male^23^ and a 76 year old female.^24^ A genetic cause of CANVAS is highly plausible given the observation of 13 affected siblings and families with multiple affected individuals over several generations.^12^ The pattern of inheritance suggests an autosomal recessive trait, although autosomal dominant inheritance with incomplete penetrance cannot be excluded. CANVAS symptoms overlap considerably with SCA3 (also known as Machado-Joseph disease, MIM: 109150) and Friedreich ataxia (MIM: 229300), both genetic forms of ataxia caused by the inheritance of a pathogenic RE. These observations are consistent with the hypothesis that a novel pathogenic STR expansion may underlie CANVAS.

Historically, the detection of REs has been time-consuming and expensive. Indeed, it is only in recent years that computational methods have been developed to screen for RE in short-read whole exome sequence (WES) and WGS data^25^, leading to the discovery of novel, disease causing REs. For example, a pentanucleotide RE was identified to underlie autosomal dominant spinocerebellar ataxia 37 (SCA37; OMIM: 615945).^26^ Moreover, pathogenic REs of intronic pentamers (TTTCA)_n_ and (TTTTA)_n_ were identified as the cause of Benign Adult Familial Myoclonus Epilepsy locus 1, 6 and 7 (BAFME1, OMIM: 618073; BAFME6, OMIM: 618074; BAFME1, OMIM: 618075).^27^

A number of bioinformatics tools now exist that allow screening of short-read sequencing data for expanded STRs.^25^ Initially, STR detection tools, such as lobSTR and hipSTR, were limited to short STRs that were encompassed by a single sequencing read. However, in the last two years, multiple methods have been released that can screen WES and WGS datasets for REs without being limited by read length. These include ExpansionHunter (EH)^28^, exSTRa^8^, TREDPARSE^29^, STRetch^30^ and GangSTR.^31^ These are all reference based methods - i.e. they rely on a catalogue of STR loci and motifs and are therefore limited to detecting expansion of previously defined STRs, such as those catalogued in the UCSC track. Moreover, the normal variability in STR length and repeat composition remains poorly described, particularly for rare STRs or those larger than ∼100bp. Therefore, there is a need for bioinformatics tools that are unbiased to the limited catalogues of STR loci available. Ideally, these tools will be able to search genome-wide for expanded repeat sequences in NGS data, independent of prior knowledge of either the location or composition of the RE. Here, we utilized a STR reference-free method called Expansion Hunter De Novo (EHdn), in combination with multiple reference-based tools, to show that CANVAS is caused by the homozygous inheritance of a novel and expanded intronic pentamer [(AAGGG)_exp]_ in the gene encoding Replication Factor C Subunit 1 (*RFC1*). An independent study, published while this work was under review, similarly identified the causal pentamer in *RFC1*. Cortese and colleagues defined a small linkage region from ten families with CANVAS and the causative RE was identified by WGS and visual inspection of the aligned read pairs inside the linkage region.^32^

## MATERIALS AND METHODS

### Recruitment, linkage and next generation sequence data

The Royal Children’s Hospital Human Research Ethics Committee approved the study (HREC 28097). Informed consent was obtained from all participants and clinical details were collected from clinical assessments and review of medical records. Genomic DNA was isolated from peripheral blood. Single nucleotide polymorphism (SNP) genotype data were generated for two affected siblings from three families (CANVAS1, 2, 3) and all six siblings from family CANVAS4 using the Illumina Infinium HumanOmniExpress BeadChip genotyping array. SNP genotypes for individuals from CANVAS9 were extracted from WES data.^33^ Parametric multipoint linkage analysis was subsequently performed using LINKDATAGEN and MERLIN^34;35^ specifying a rare recessive disease model with complete penetrance, and overlapping linkage signals were detected using BEDtools.^36^ WES was performed on individuals from CANVAS9 using Agilent SureSelect XT Human All exon V5 + UTR on the Illumina HiSeq2000 platform at 50x mean coverage. WES was performed on an additional 23 individuals from 15 families in collaboration with the Johns Hopkins Center for Inherited Disease Research (CIDR) as part of the Baylor-Hopkins Center for Mendelian Genomics (BHCMG). WGS was performed in two stages. Libraries for the first round of samples, including two affected individuals from CANVAS1 and CANVAS9 and 31 individuals lacking a clinical diagnosis of CANVAS (subsequently referred to as controls although some have a diagnosis other than CANVAS), were prepared using the TruSeq nano PCR-based Library Preparation Kit and sequenced on the Illumina HiSeq X platform. Libraries for the second round of WGS, including affected individuals with evidence of an alternate RE motif (CANVAS2 and CANVAS8) or lacking the pathogenic RE in *RFC1* (CANVAS11,13, 17 and 19), were prepared using the TruSeq PCR-free DNA HT Library Preparation Kit and sequenced on the Illumina NovaSeq 6000 platform. PCR-free WGS data from 69 unrelated Coriell controls^28^ was obtained from Illumina. GTEx samples (SRA files, 133 WGS with matching cerebellar RNA-seq) were downloaded from the dbGAP (phs000424.v7.p2).

### Alignment and variant calling

Alignment and haplotype calling were performed based on the GATK best practice pipeline. All WES and WGS datasets were aligned to the hg19 reference genome using BWA-mem, then duplicate marking, local realignment and recalibration were performed with GATK. Merged VCF files were annotated using vcfanno^37^ and ANNOVAR.^38^ Candidate variant filtering was performed using CAVALIER, an R package for variant interpretation in NGS data (https://github.com/bahlolab/cavalier). Standard variant calling was performed on WGS data for CANVAS samples negative for the pathogenic RE in *RFC1*. Candidate variants were defined as i) occurring in known ataxia genes, as defined by OMIM, ii) exonic, with a minor allele frequency of less than 0.0001 in gnomAD (both genome and exome data) and iii) predicted pathogenic by both SIFT and PolyPhen2. RNA-seq data was aligned to the hg19 reference genome (ENSEMBL Homo_sapiens.GRCh37.75) using STAR.^39^ Reads were summarized by gene ID into a counts matrix using featureCounts^40^ (quality score >= 10) and converted to log10 of the counts per million using limma.^41^

### STR analysis

Genome-wide screening for putative REs was performed using Expansion Hunter Denovo (EHdn) version 0.6.2, an open-source method that is being developed by Illumina, Inc, the Walter Eliza Hall Institute and others. EHdn operates by performing a genome-wide search for read pairs where one mate has confident alignment (anchor) and the second mate consists of repetition of a repeat motif (in-repeat read). The program reports the counts of in-repeat reads with anchor mates stratified by the repeat motif and genomic position of their anchor mate. For this analysis we defined a confidently-aligned read as one aligned with MAPQ of 50 or above. The counts of in-repeat reads with anchor mates were subsequently compared for each region in cases (CANVAS) and controls using a permutation test (10^^^6 permutations). The resulting p-values were used to rank candidate sites with higher counts in individuals with CANVAS than in the controls for further computational validation. These candidates were subsequently annotated with ANNOVAR.

Computational validation was performed using five independent STR detection tools for short-read NGS after updating the STR catalogue reference files to incorporate the identified motifs. The RE candidates were screened in the two individuals with CANVAS and the 31 non-CANVAS controls using exSTRa and EH, then the top candidate [(AAGGG)_exp_ STR in *RFC1*] was further validated with TREDPARSE, GangSTR and STRetch. All tools were used with default parameters, with the following additional parameters for EH: read-depth of 30 and min-anchor-mapq of 20. All five tools were also used to screen for the (AAGGG)_exp_ *RFC1* STR in the 69 Coriell control WGS datasets. A short-list of (AAGGG)_exp_ carriers was generated based on consensus calling from at least four of the five tools.

Individuals diagnosed with CANVAS lacking the (AAGGG)_exp_ *RFC1* RE were further screened with EHdn for novel STRs and for known pathogenic STRs using exSTRa and EH. The WES datasets could not be analyzed for the (AAGGG)_exp_ *RFC1* RE as the intronic locus (chr4:39350045-39350095, hg19) was not captured during library preparation. However, the region was visualized using the Integrative Genomics Viewer (IGV) to identify potential off-target reads which could provide supportive evidence for the presence of the (AAGGG)_exp_ motif. Only samples with at least one read mapping at the STR in the *RFC1* locus were considered.

### Haplotyping and mutation dating

Haplotyping was performed on the WES data. Variants were filtered based on read depth (≥30), including both exonic and non-exonic variants. A core haplotype was defined based on sharing amongst a majority of affected individuals. A method based on haplotype sharing^42^ was used to determine the most recent common ancestor (MRCA) from whom the core haplotype was inherited, as well as dating additional sub-haplotypes shared by clusters of individuals, which are likely to be individuals with a MRCA who is more recent than that for the whole group (https://shiny.wehi.edu.au/rafehi.h/mutation-dating/).

### Molecular genetic studies

We designed a PCR assay to test for presence of additional inserted sequence, not present in the reference database, at the *RFC1* STR. The primers (Table S1) flank the STR and are predicted to amplify a 253bp fragment using standard PCR conditions with a 30 second extension cycle. Presence of the pathogenic *RFC1* RE was tested by repeat-primed PCR utilizing three primers; TPP_CANVAS_FAM_2F, 5R_TPP_M13R_CANVAS_RE_R and TPP_M13R (Table S1). The FAM labelled forward primer is locus specific, while the repeat-specific primer (5R_TPP) includes a tag M13R sequence. PCR was performed in a 20µl reaction with 20 ng genomic DNA, 0.8 µM of both the FAM labelled forward primer and TPP_M13R and 0.2 µM 5R_TPP using GoTaq^®^ Long PCR Polymerase (Promega). A standard 60TD55 protocol was utilized (94°C denaturation for 30 s, 60TD55°C anneal for 30 s, and 72°C extension for 2 min), products were detected on an ABI3730xl DNA Analyzer and visualized using PeakScanner 2 (Applied Biosystems).

## RESULTS

### Case recruitment

The workflow for this study is summarized in Figure 1. Individuals with a clinical diagnosis of CANVAS were recruited following neurological assessment and investigation in accordance with published guidelines.^43^ While variable between cases, data leading to the clinical diagnosis included evidence of combined cerebellar and bilateral vestibular impairment, cerebellar atrophy on MRI, neurophysiological evidence of impaired sensory nerve function and negative genetic testing for pathogenic RE at common SCA loci (typically *SCA1, 2, 3, 6* and *7*) and *FRDA* (Friedreich ataxia, FRDA). In total, the cohort consisted of 35 individuals with a clinical diagnosis of CANVAS (Table 1). The individuals came from eleven families with a single affected individual, seven families with affected sib pairs and four larger/multigenerational families (Figure S1). A full clinical description of the cohort will be reported in a forthcoming manuscript.

**Table 1:** Clinical features and genetic analysis of *RFC1* locus in study participants.

| Family | Participants (sex) | SNP array | WES | WGS | *RFC1* STR in WES | PCR wildtype allele | Repeat-primed PCR | Genetic Diagnosis | Haplotype | Ethnicity |
| --- | --- | --- | --- | --- | --- | --- | --- | --- | --- | --- |
| CANVAS1 | 2 (F) | ✓ | ✓ | ✓ | ND | ✗ | ✓ | CANVAS | A/other | European |
| CANVAS2 | 2 (M) | ✓ | ✓ | ✓ | AAGGG and AAAGG | ✗ | ✓ | CANVAS | A | European |
| CANVAS3 | 2 (F) | ✓ | ✓ | ✗ | AAGGG | ✗ | ✓ | CANVAS | A | European |
| CANVAS4 | 4 (3M, 1F) | ✓ | ✓ | ✗ | AAGGG | ✗ | ✓ | CANVAS | A | Greek-Cypriot |
| CANVAS5 | 2 (M, F) | ✗ | ✗ | ✗ | ND | ✗ | ✓ | CANVAS | Not assessed | Not reported |
| CANVAS6 | 2 (M) | ✗ | ✓ | ✗ | AAGGG | ✗ | ✓ | CANVAS | A | Lithuanian/Latvian |
| CANVAS7 | 1 (M) | ✗ | ✓ | ✗ | ND | ✗ | ✓ | CANVAS | A | European-Maori |
| CANVAS8 | 1 (F) | ✗ | ✓ | ✓ | AAGGG and AAAGG | ✗ | ✓ | CANVAS | A/other | European |
| CANVAS9 | 4 (1M, 3F) | ✗ | ✓ | ✓ | AAGGG | ✗ | ✓ | CANVAS | A | Lebanese |
| CANVAS10 | 1 (M) | ✗ | ✓ | ✗ | AAGGG | ✗ | ✓ | CANVAS | A | European |
| CANVAS11 | 1 (M) | ✗ | ✓ | ✓ | ND | ✓ | ✗ | ? | NA | Anglo-saxon |
| CANVAS12 | 1 (M) | ✗ | ✓ | ✗ | ND | ✗ | ✓ | CANVAS | A | Turkish |
| CANVAS13 | 1 (M) | ✗ | ✓ | ✓ | Reference | ✓ | ✗ | SCA3 | NA | Martinique |
| CANVAS14 | 1 (M) | ✗ | ✓ | ✗ | AAGGG | ✗ | ✓ | CANVAS | Other* | European |
| CANVAS16 | 1 (F) | ✗ | ✗ | ✗ | NA | ✗ | ✓ | CANVAS | Not assessed | European |
| CANVAS17 | 2 (M) | ✗ | ✓ | ✓ | Reference | ✓ | ✗ | SACS | NA | European |
| CANVAS18 | 1 (F) | ✖ | ✓ | ✖ | ND | ✖ | ✓ | CANVAS | A | European-Maori |
| CANVAS19 | 1 (F) | ✖ | ✖ | ✓ | NA | ✓ | ✖ | SCA45 | Not assessed | European |
| CANVAS20 | 2 (1M, 1F) | ✖ | ✖ | ✖ | NA | ✖ | ✓ | CANVAS | Not assessed | Spanish |
| CANVAS21 | 1 (M) | ✖ | ✖ | ✖ | NA | ✖ | ✓ | CANVAS | Not assessed | Indian |
| CANVAS22 | 1 (M) | ✖ | ✖ | ✖ | NA | ✖ | ✓ | CANVAS | Not assessed | Hungarian |
| CANVAS23 | 1 (U) | ✖ | ✖ | ✖ | NA | ✖ | ✓ | CANVAS | Not assessed | Not reported |

M=male, F=female, U=deidentified, NA=not applicable, ND=not detected, Other\*= a different haplotype OR shortened A haplotpye
The gene reference sequences utilized were NC\_000004 and NM\_002913 (RFC1).

**Figure 1:**
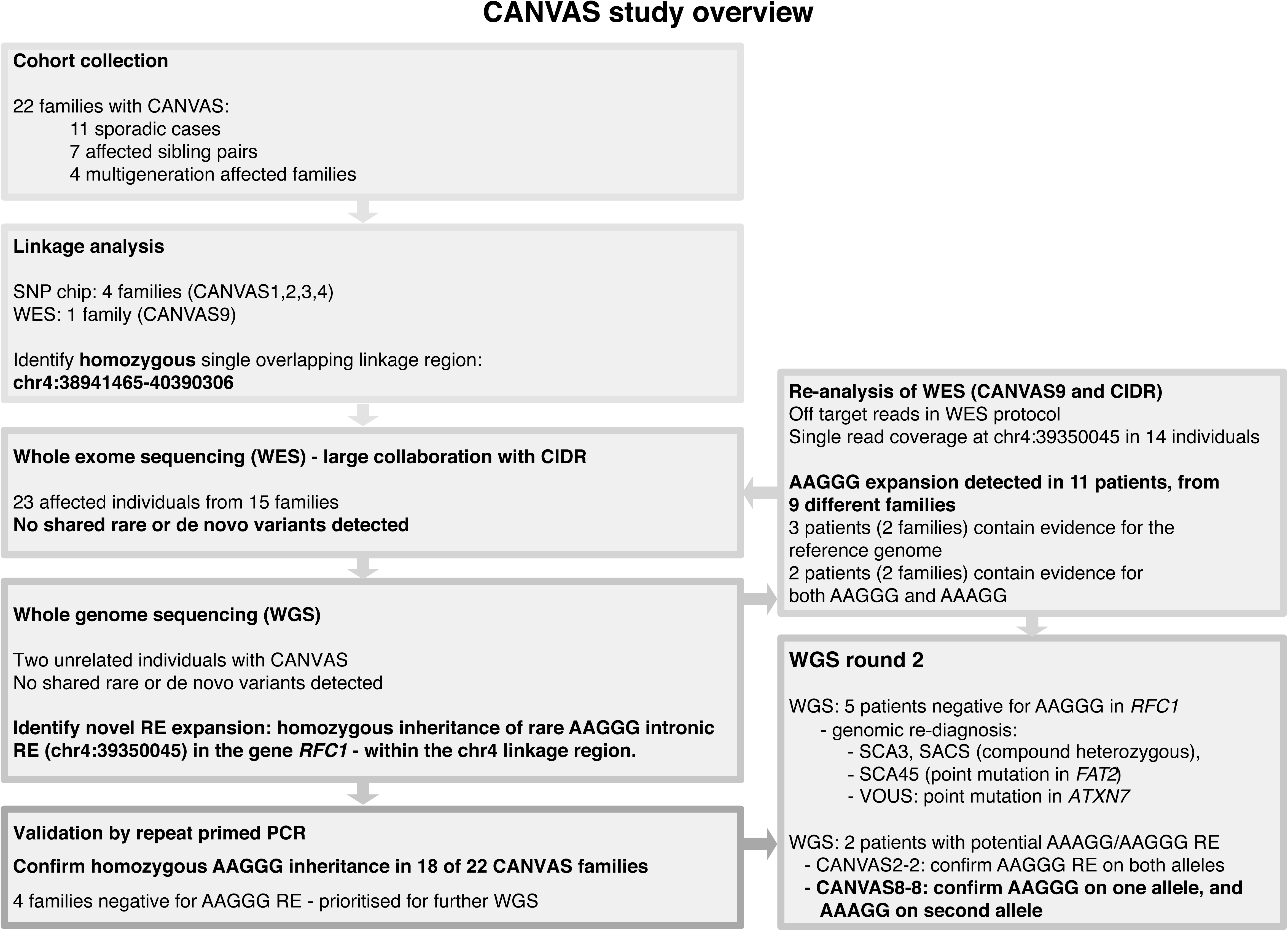
Overview of the CANVAS study and genetic investigations performed.

### Linkage analysis

CANVAS typically presents in families with one or multiple affected individuals in a single generation, consistent with a recessive inheritance. For example, in the second-degree consanguineous family CANVAS9, four siblings were diagnosed with CANVAS and two were classified as unaffected at the time phenotyping was performed (Figure 2A). Parametric multipoint linkage analysis was performed on five CANVAS families (CANVAS1, −2, −3, −4 and −9, Figure S1) specifying a rare recessive disease model with complete penetrance. This identified linkage regions with logarithm of odds (LOD) scores ranging from 0.6 for smaller pedigrees (two affected siblings), to a statistically significant linkage region on chromosome 4 in CANVAS9 (LOD=3.25, Figure 2B). Intersection of the linkage regions from the five families identified a single region on chromosome four (chr4:38887351-40463592, hg19, combined LOD=7.04) common to all families (Figure 2C). CNV analysis utilizing PennCNV did not identify any potential copy number variants in the minimal linkage region.^44^ The 1.5MB shared region contains 42 genes, of which 14 are protein coding, none with any association with ataxia in OMIM or the published literature (Table S2).

**Figure 2:**
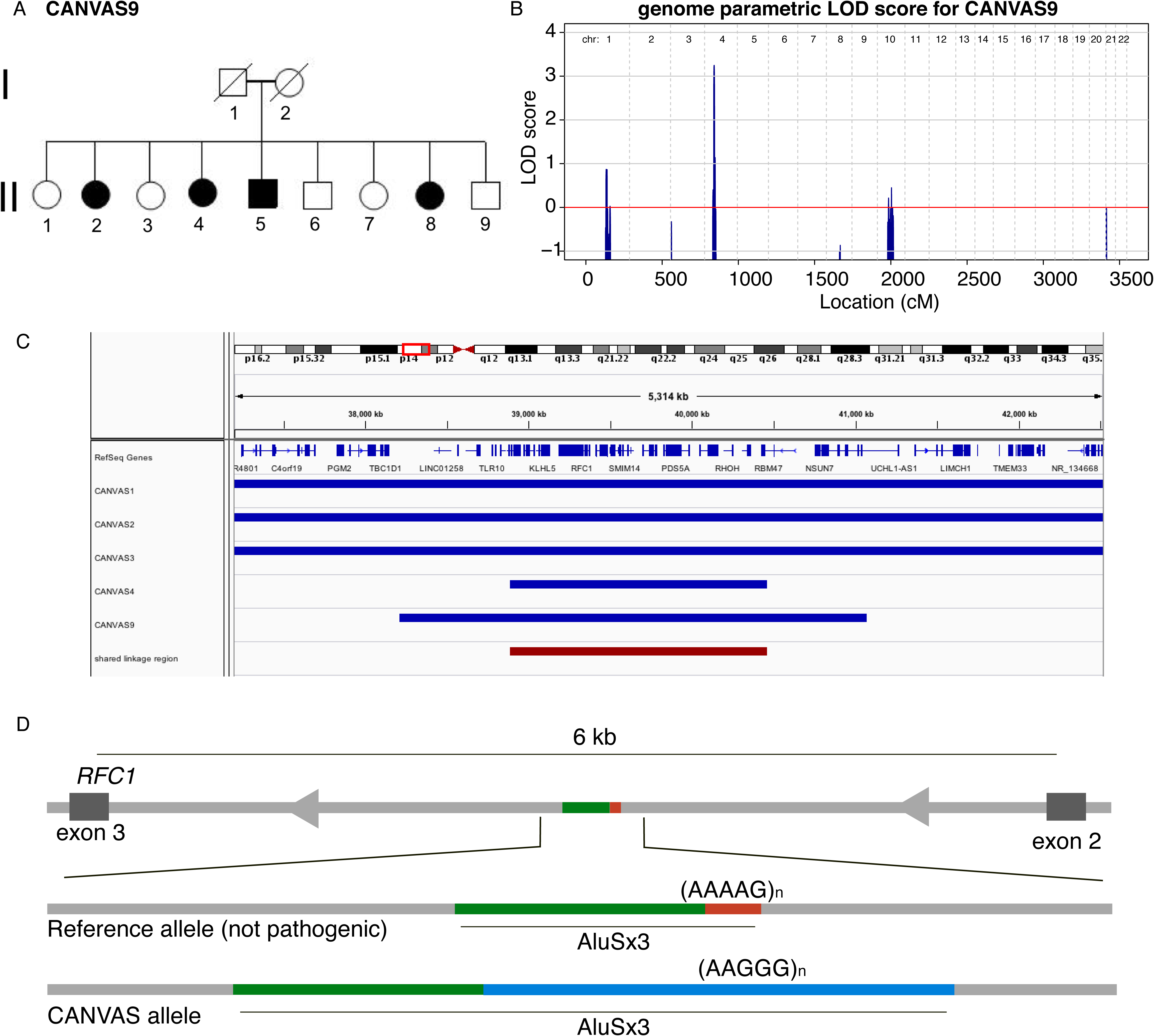
Linkage of the CANVAS locus to chromosome 4 and identification of (AAGGG)_exp_ intronic insertion in *RFC1*. A. The pedigree of the family CANVAS9 highlights the apparent recessive inheritance pattern. B. Linkage analysis of CANVAS9 identified significant linkage to chromosome 4 (LOD=3.25). C. Linkage regions for individual families CANVAS1, 2, 3, 4 and 9 are shown in blue and the overlapping region shown in red (chr4:38887351-40463592, combined LOD=7.04). D. STR analysis of WGS from two unrelated individuals with CANVAS identified a novel expanded STR in the second intron of *RFC1*. The (AAAAG)_11_ motif that is present in the reference genome and part of an existing Alu element (AluSx3) is replaced by the (AAGGG)_exp_ RE.

### Large-scale WES analysis did not identify candidate pathogenic variants

WES was used to screen 27 affected individuals with CANVAS from 15 families for potentially pathogenic rare variants (MAF < 0.001) shared across multiple pedigrees in a homozygous or compound heterozygous inheritance pattern. No candidate mutations were detected, either within the chromosome 4 linkage region or elsewhere in the genome.

### Identification of a novel (AAGGG)_exp_ RE in the linkage region

The lack of candidate variants identified from the WES data suggested the possibility of (i) intronic or intergenic mutations, or (ii) that CANVAS might be caused by a non-standard mutation, such as a pathogenic RE of an STR. Therefore, WGS was performed on two individuals from different pedigrees (CANVAS1 and CANVAS9) who share the chr4 linkage region. EHdn was used to perform a genome-wide screen for STRs in the two individuals with CANVAS compared to WGS data from 31 unrelated controls. This identified 19 regions with a p value<0.005 (Table S3), although genome-wide significance could not be achieved after adjustment for multiple testing due to the skewed ratio of the number of cases to controls (2 versus 31). These candidate STRs were visualized with the Integrative Genomics Viewer (IGV) tool, which suggested that the (AAGGG)_exp_ STR within intron 2 of the gene encoding Replication Factor C1 (*RFC1*) was likely real and present in both alleles in the affected individuals, consistent with the recessive inheritance pattern hypothesized for CANVAS (Figure S2). In addition, this was the only candidate that (I) was localized to the chr4 linkage region and (II) was able to be validated using existing STR detection tools (see below). In both individuals with CANVAS, the novel (AAGGG)_exp_ pentamer replaced an (AAAAG)_11_ motif located at the same position in the reference genome (chr4:39350045-39350095, hg19) and appeared to be significantly expanded compared to controls. Visualization of the region in the UCSC genome browser identified that the reference motif (AAAAG)_11_ is the 3’ end of an Alu element, AluSx3. In individuals with CANVAS, the (AAAAG)_11_ motif is substituted by the (AAGGG)_exp_ motif, with potential interruptions to the Alu element (Figure 2D).

### Confirmation of (AAGGG)_exp_ STR in off-target WES reads

While WGS was only performed in two individuals with CANVAS, the majority of the cohort (n=27) was analyzed by WES. The putative pathogenic CANVAS RE is located in intron 2 of *RFC1*, 2863bp downstream of exon 2 and 2952bp upstream of exon 3. Therefore WES data is *a priori* assumed to be uninformative for this RE as it is not targeted during DNA capture. However, given that WES data includes off-target reads, we hypothesized that some reads might map to the *RFC1* RE locus. Visual assessment of the WES data in IGV identified 14 individuals with informative reads; the maximum off-target read coverage at this locus was two, with a median of one read. While three individuals only had reads that correspond to the reference genome STR sequence (AAAAG), eleven affected individuals from nine families had reads containing (AAGGG) repeats (Table 1). Furthermore, single affected individuals from families CANVAS2 and CANVAS8 had single, independent reads identifying (AAGGG) and (AAAGG) motifs at the *RFC1* STR locus. This observation raised the possibility that CANVAS might result from pathogenic expansions of different pentanucleotide motifs.

### Computational validation with existing STR detection tools

Multiple tools have been developed in recent years that test for the presence of REs at pre-defined STRs. Therefore, we inserted the novel *RFC1* STR motifs into the STR reference files and used exSTRa, EH, TREDPARSE, STRetch and GangSTR to estimate the size of the STR, and/or detect REs in the WGS data from the two original CANVAS samples (CANVAS1 and 9) and seven additional individuals with CANVAS. The seven additional CANVAS samples selected for WGS were those with WES evidence for an alternate (AAAGG) motif (families CANVAS2 and 8), and those who did not appear to have a RE at the *RFC1* locus based on the PCR/RP-PCR studies described below (CANVAS11, 13, 17 and 19 families). The library preparation this second round of WGS was PCR-free as PCR amplification has previously been shown to affect RE detection.^8^ Using exSTRa, we confirmed the homozygous inheritance of the (AAGGG)_exp_ RE in three individuals (CANVAS1, 2 and 9, Figure 3). The empirical cumulative distribution function (ECDF) pattern for CANVAS2 is consistent with the presence of one shorter and one longer (AAGGG)_exp_ RE, while CANVAS8 appears to only have a single (AAGGG)_exp_ allele. Screening of all datasets for the (AAAGG)_exp_ motif at the chr4 *RFC1* locus using exSTRa identified an expansion of this STR only in CANVAS8, suggesting this individual is a compound heterozygote, with an (AAGGG)_exp_ RE on one allele and an (AAAGG)_exp_ RE on the other. Visualization in IGV confirmed the presence of the (AAAGG)_n_ motif embedded within the reference STR: (AAAAG)_6_-(AAAGG)_n_-(AAAAG)_6_ (Figure S3). This observation raises the possibility that an expanded (AAAGG)_exp_ motif might also be associated with CANVAS.

**Figure 3:**
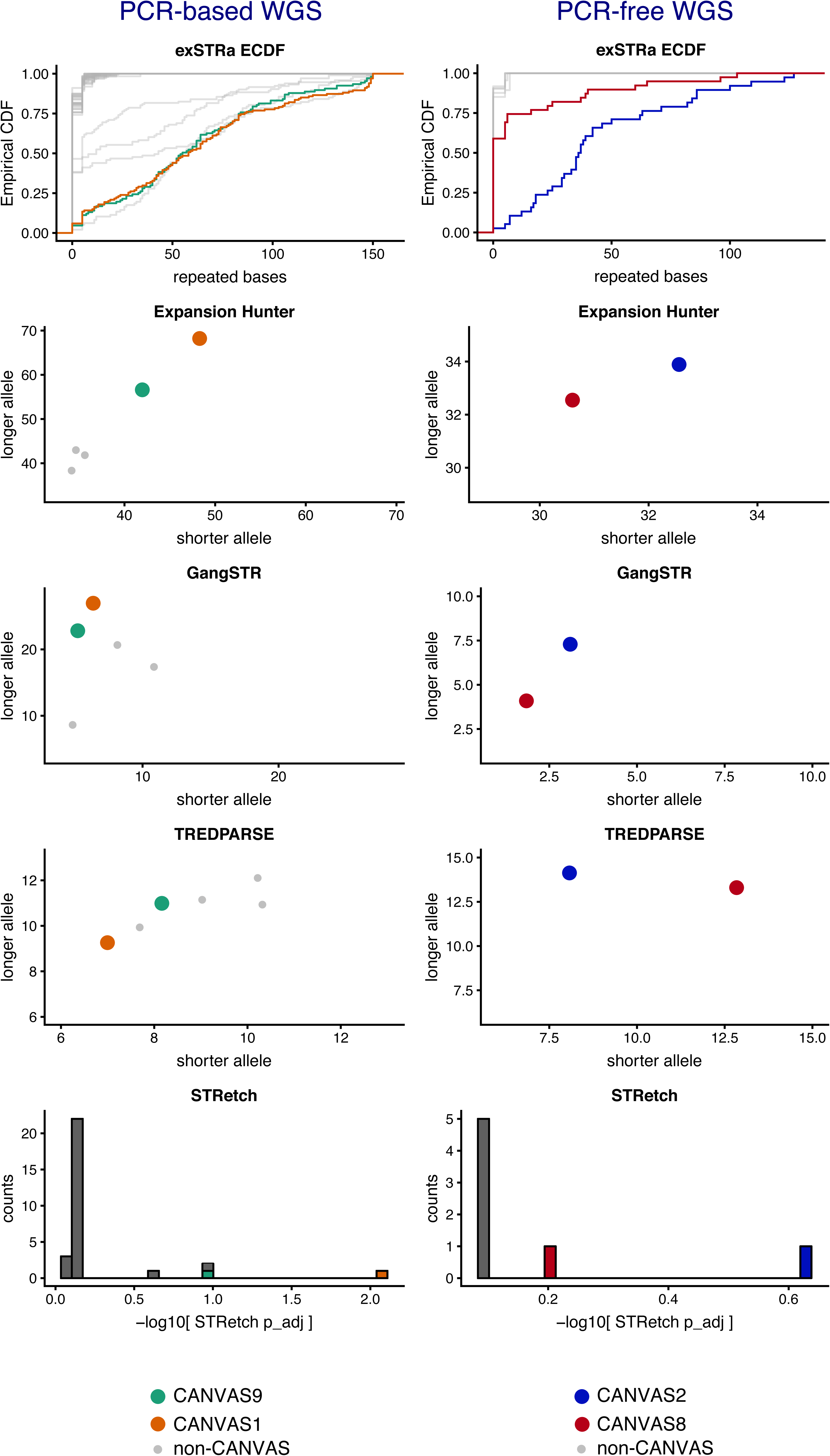
Computational validation of the (AAGGG)_exp_ RE. The (AAGGG)_exp_ RE at the coordinates chr4:39350045-39350095 was added to the reference databases of the tools exSTRa, EH, GangSTR, TREDPARSE and STRetch and WGS data from four unrelated individuals with CANVAS was analysed [CANVAS1 (orange), CANVAS2 (blue), CANVAS8 (red) and CANVAS9 (green)]. The non-CANVAS controls are presented in grey. Plots have been divided into PCR-based and PCR-free WGS (left and right columns, respectively). The Y and X axes for ExpansionHunter, GangSTR and TREDPARSE refer to the number of repeat units on the longer and shorter allele per individual, respectively. The Y axis for the STRetch plot refers to the number of individuals.

EH, TREDPARSE and GangSTR were used to estimate the length of the AAGGG motif on each allele (Figure 3). The results were highly variable depending on the tool used. EH reported minimum and maximum allele sizes ranging from 30 to 68 in individuals with the RE. The allele size ranges estimated by GangSTR and TREPARSE were 2 to 27 and 7 to 14, respectively. Furthermore, all three tools inferred the presence of two alleles, even in individuals who carry a single allele, and hence do not appear to be distinguishing read contributions between the alleles, also contributing to unreliable size estimates. Reads comprised of the (AAGGG)_exp_ motif in particular also showed evidence of high read sequencing error. Based on these results, we can infer that while the CANVAS samples were all correctly identified as having homozygous RE at *RFC1*, estimates of expansion size are inconsistent and appear likely to significantly underestimate the actual repeat size.

The consensus of the different tools was that CANVAS11, 13, 17 and 19 families did not encode a pathogenic RE [either (AAGGG)_exp_ or (AAAGG)_exp_] at the *RFC1* locus, which we confirmed by PCR analyses (see below). However, the *RFC1* (AAGGG)_exp_ RE was present in three of the control WGS datasets [two heterozygous, one homozygous, allele frequency ∼0.06 (4/62), Figure 3]. No control individuals were identified to carry the (AAAGG)_exp_ motif. As with the CANVAS samples, the STR sizing estimates using the different tools was inconsistent, therefore no conclusions could be drawn from this *in silico* analysis regarding the relative size of the (AAGGG)_exp_ RE in controls compared to individuals with CANVAS. We then analyzed a larger in-house collection of unrelated control Coriell WGS samples (N=69) and again failed to identify the (AAAGG)_exp_ motif. However, we identified six individuals heterozygous for the (AAGGG)_exp_ RE, representing a frequency estimate of ∼0.04 [(6/138) (Figure S4)]. Using the NGS QC software tool peddy, we found evidence that two of these heterozygous individuals are of European ancestry and that two further individuals are of admixed Native American ancestry. Finally, we accessed WGS from GTEx for 133 individuals who have matching brain (cerebellum) RNA-seq. Our analysis identified 11 heterozygous carriers of the (AAGGG)_exp_ RE, representing an estimated allele frequency of ∼0.04 (11/266), consistent with our in-house collection.

### Validation of the (AAGGG)_n_ RE as the causal variant for CANVAS

We developed a PCR assay that amplifies across the repeat tract to rapidly screen for the presence of a non-expanded allele at the *RFC1* STR. Although the screen does not distinguish between the (AAAAG)_11_ reference STR, the (AAGGG) STR, or any other potential motif, amplification of an ∼250bp fragment indicates at least one allele is not expanded. Moreover, the presence of two distinct non-expanded products is indicative of a heterozygous non-mutant state. Conversely, the complete absence of the PCR product provides indirect evidence of a RE affecting both alleles of the *RFC1* locus. Analysis of all available DNA samples from individuals with CANVAS suggested that the reference STR at the *RFC1* locus was not present in 30 clinically diagnosed individuals from 18 (of 22) CANVAS families (Table 1, Figure 4). Notably, unaffected individuals from *RFC1* positive families carried at least one non-expanded *RFC1* allele (Figure S5). To directly confirm expansion of the novel (AAGGG) motif in *RFC1*, we developed a locus specific repeat-primed PCR assay, using a primer located adjacent to the *RFC1* repeat and an AAGGG-specific primer. Consistent with the PCR assay, affected individuals from the 18 families demonstrated a saw-toothed ‘ladder’ when the repeat-primed PCR products were analyzed by capillary array (Table 1, Figure 4). These results suggest a homozygous RE underlies CANVAS in these 18 families, and at least one pathogenic allele encodes the (AAGGG)_exp_ RE. Molecular analysis of the DNA for the three in-house control individuals with the *in silico* predicted (AAGGG)_exp_ motif (Figure 3) demonstrated a ∼250bp product in both heterozygous samples but no product in the homozygous individual. The repeat-primed assay demonstrated a saw-toothed ladder in all three samples (Figure S6). Collectively, these analyses suggested all three control individuals have at least one copy of the pathogenic (AAGGG)_exp_ RE at *RFC1*, although the size of the RE cannot be determined by these analyses.

**Figure 4:**
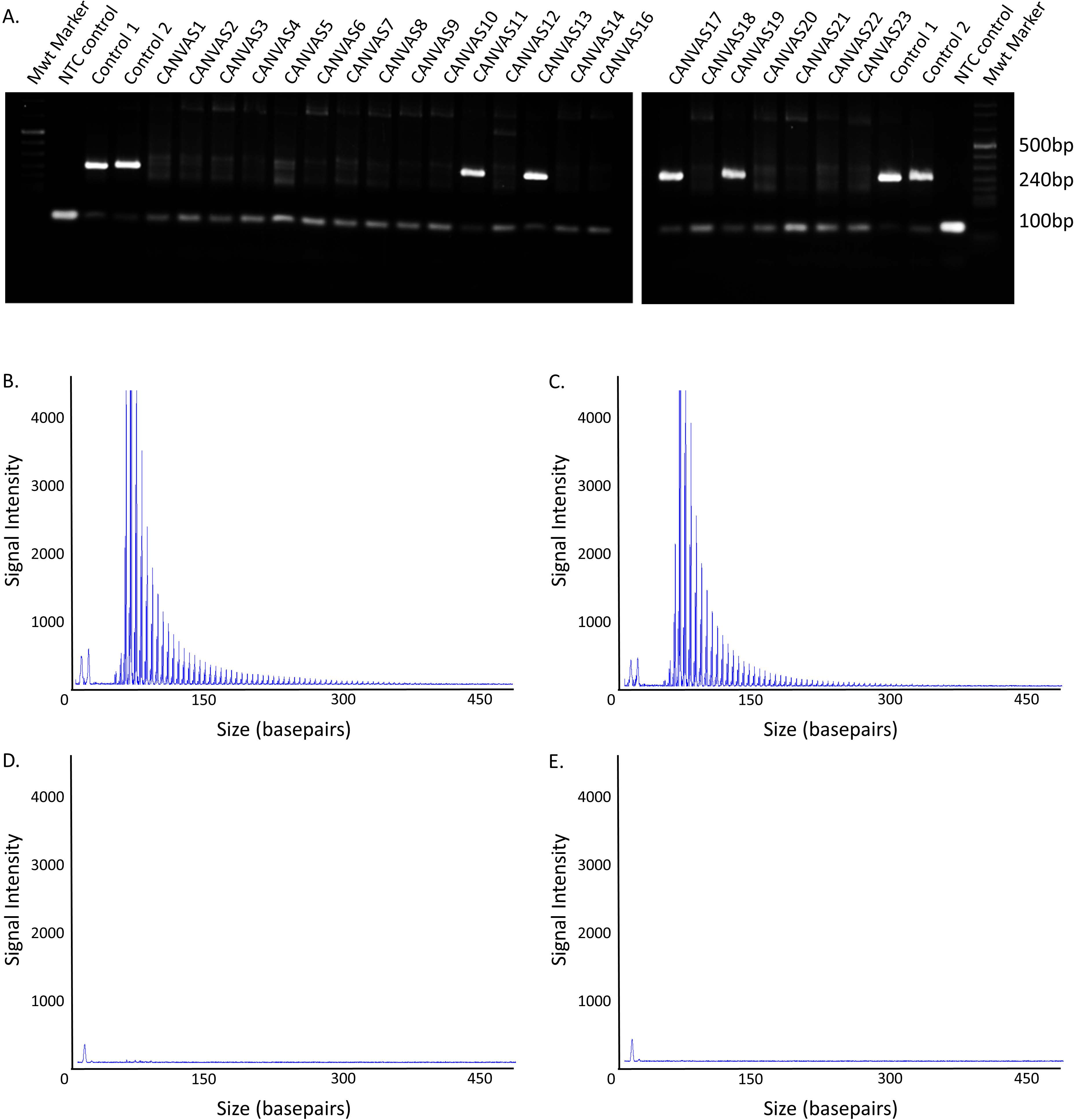
Genetic validation of the (AAGGG)_exp_ RE. A. PCR analysis of the *RFC1* STR failed to produce the control ∼253bp reference product in 18 of 22 CANVAS families. Representative images of the repeat-primed PCR for the (AAGGG)_exp_ RE demonstrating a saw-toothed product with 5 base pair repeat unit size, amplified from gDNA of individuals from CANVAS1 (B) and CANVAS9 (C). No product was observed for the unaffected control (D) and no gDNA template negative control (E).

In four families (CANVAS11, 13, 17 and 19) the presence of the expected reference PCR amplicon and lack of a repeat-primed PCR product suggested the pathogenic *RFC1* RE was not present on either allele. This implied that these individuals have a different CANVAS-causing mutation in *RFC1*, or there is locus heterogeneity. A third possibility is that they do not have CANVAS but instead a related ataxia. Therefore, we performed WGS on these individuals and initially screened for known REs associated with ataxias using exSTRa and EH. A CAG trinucleotide expansion in *ATXN3*, associated with spino-cerebellar ataxia type 3 (SCA3; OMIM 109150 also known as Machado-Josephs disease) was identified in CANVAS13 (Figure S7) and confirmed by diagnostic testing. The WGS was then screened for novel or rare SNPs and indels in genes known to cause ataxia. No *de novo* or rare variants were identified in *RFC1* however a potential genomic re-diagnosis was achieved in two additional families. In CANVAS17 two variants [NM_001278055:c.12398delT, p.(Phe4133Serfs*28) and NM_001278055:c.5306T>A, p.(Val1769Asp)] were identified in the gene encoding sacsin (*SACS*) and segregation analysis confirmed they were in trans. Biallelic mutations in *SACS* cause spastic ataxia of the Charlevoix-Saguenay type (MIM: 270550). In CANVAS19, a heterozygous variant in the gene encoding FAT tumor suppressor homolog 2 [*FAT2*, NM_001447.2:c.4370T>C, p.(Val1457Ala)] was identified. Heterozygous mutations in *FAT2* have recently been associated with SCA45 (MIM: 604269).^45^ No potentially pathogenic variants were identified in CANVAS11, however a variant of unknown significance was identified in the gene encoding Ataxin 7 [*ATXN7*, NM_001177387.1:c.2827C>G, p.(Arg943Gly)]. CANVAS11 was also screened genome-wide with EHdn for potentially pathogenic novel RE, however no additional candidate REs were identified.

### A single founder event for the (AAGGG)_n_ RE in *RFC1*

We performed haplotype analysis to determine if the (AAGGG)_exp_ RE arose more than once in human history. Analysis of haplotypes inferred from the WES data identified a core ancestral haplotype, comprised of 27 SNPs (Figure 5A), that was shared by most individuals except CANVAS14 (Table 1, Table S4). The core haplotype spans four genes (*TMEM156, KLHL5, WDR19* and *RFC1*) and is 0.36 MB in size (chr4:38995374-39353137 (hg19)). Inspection of this region in the UCSC browser suggested that the core haplotype overlaps with a region of strong linkage disequilibrium in European and Asian populations (Han Chinese and Japanese from Tokyo), but not the Yoruba population (an ethnic group from West Africa, Figure 5B). Using a DNA recombination and haplotype-based mutation dating technique^42^, we estimate that the most recent common ancestor (MRCA) of the CANVAS cohort lived approximately 25,880 (CI: 14080-48020) years ago (Figure 5C). This age estimate corresponds to the size of the haplotype and LD block and is roughly equivalent to the origin of modern Europeans as represented by the HAPMAP CEU cohort. Further investigation of the haplotypes allowed us to infer a simple phylogeny based on identified clusters of shared haplotypes extending beyond the core haplotype, suggesting that some individuals have common ancestors more recent than that of the MRCA for the whole group. This approach identified four subgroups. Group A had a MRCA dating back 5,600 (CI: 2120-15520) years and group B (further divided into groups B1 and B2) have a MRCA dating back 4,180 years (CI: 2240-7940). Furthermore, one individual shared part of their haplotype with both groups A and B, suggesting that group B is a distant branch of the MRCA of group A. Another subgroup, C, has a MRCA that lived 1860 (CI: 560-7020) years ago. The final group labelled N, do not have any additional sharing beyond the core haplotype.

**Figure 5:**
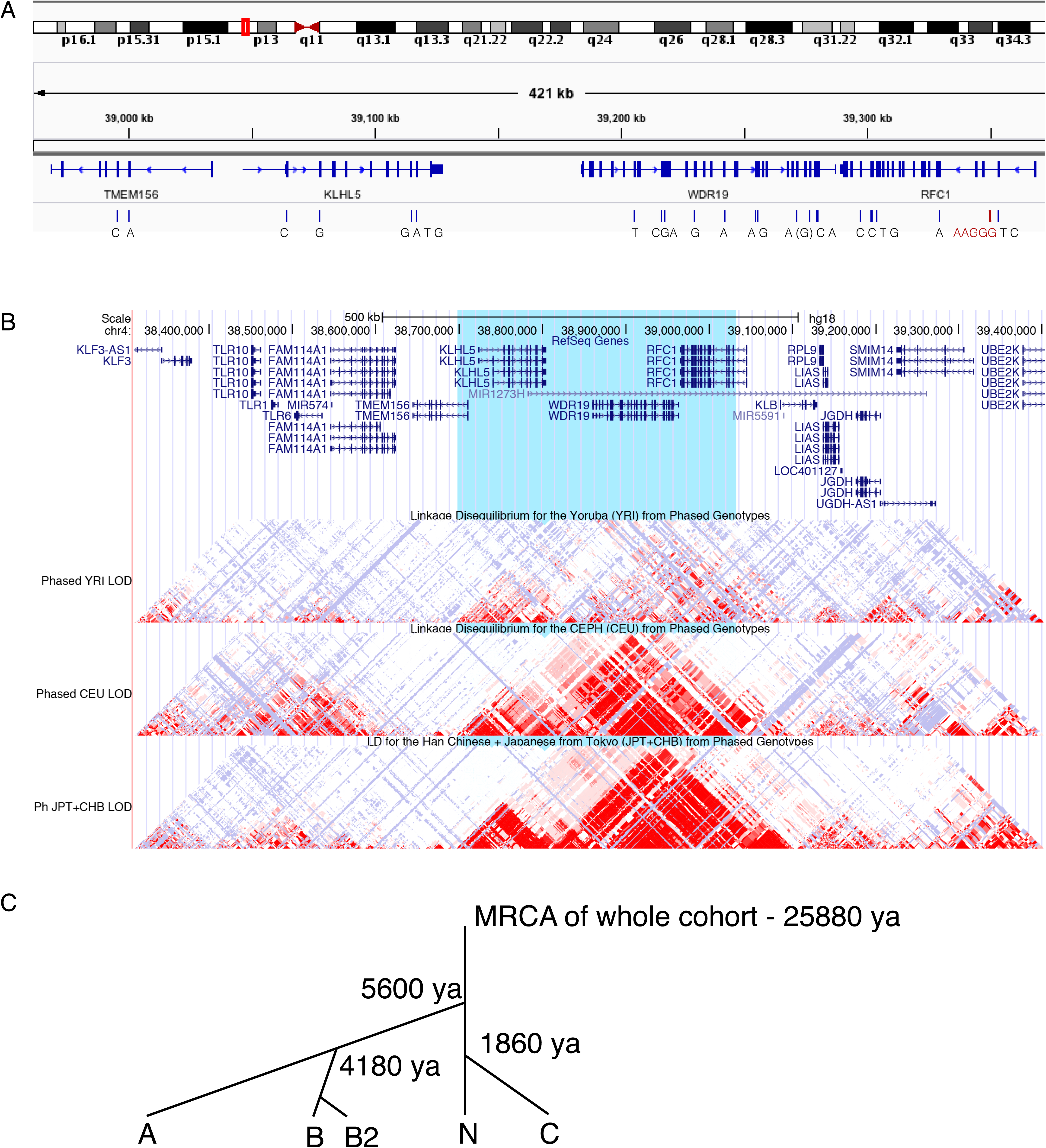
The majority of individuals with CANVAS encode an ancestral haplotype. A. Analysis of WES data identified an ancestral haplotype surrounding *RFC1* in all affected individuals confirmed to carry the (AAGGG)_exp_ RE. B. The core haplotype (blue highlight) was intersected with the linkage disequilibrium (LD) track in the UCSC browser (converted to hg18 coordinates). The three LD tracks represent the Yoruba population (top track), Europeans (middle) and Han Chinese and Japanese from Tokyo (bottom). Red areas indicate strong linkage disequilibrium. The core CANVAS haplotype spans a large LD block in Europeans, which is broken up into two LD blocks in Japanese and Chinese, suggesting an ancient origin for the CANVAS repeat expansion allele. C. Haplotype sharing between individuals with CANVAS was used to determine the age of the most recent common ancestor (MRCA) of the cohort.

Next, we compared the haplotype of the nine control samples (three in-house controls and six from the Coriell collection) that carry the (AAGGG)_exp_ RE to the core haplotype defined in the individuals with CANVAS. All controls shared at least part of the core haplotype, again suggesting that the (AAGGG)_exp_ RE arose once in history. Finally, we determined that nine of the 11 individuals from GTEx heterozygous for the (AAGGG)_exp_ RE also shared the same core haplotype identified in individuals with CANVAS. The haplotype-specific SNP rs2066782 (exon 18, chr4:39303925, A>G) enabled us to analyse the expression of the (AAGGG)_exp_ *RFC1* allele in the cerebellum RNA-seq data and confirm that the STR did not inhibit the expression of *RFC1* compared to the reference (AAAAG)_11_ allele. The remaining two carriers do not appear to share the core haplotype. As they do not have heterozygous SNPs in their exons, allele specific expression could not be determined.

## DISCUSSION

Since the first description of the syndrome of cerebellar ataxia with bilateral vestibulopathy in 2004^46^ and proposal of CANVAS as a distinct clinical entity in 2011^11^ there has been little progress made in delineating the etiology of the disorder. While most affected individuals are described as idiopathic, reports of multiple affected sib pairs^12^ and a family with three affected individuals^47^ have suggested that an autosomal recessive mode of inheritance is most likely. The genetic basis of CANVAS has now been identified and validated in two independent studies, one recently published by Cortese *et al*^32^ and this study. Both studies utilized a similar study design, with linkage analysis to reduce the genomic search space to a modest interval (<2Mb), but no plausible causal variant(s) could be identified in WES data. WGS was then performed on multiple individuals and Cortese *et al* successfully identified the RE by visual inspection of the aligned read pairs inside the linkage region using the Integrative Genomics Viewer. In contrast, we utilized a bioinformatics approach and performed genome-wide analysis of WGS data to identify potential RE and then prioritized the RE located within the linkage interval. While both approaches were successful, the bioinformatics approach to RE detection, as described in this study, is likely more sensitive and practical, and can be applied even in the absence of a small, or indeed any, linkage region. Furthermore, using a bioinformatics approach allows simultaneous testing of other potentially causal RE due to differential diagnoses. For example, we quickly re-diagnosed an affected individual with a pathogenic SCA3 RE.

Previously, the only variant associated with CANVAS was a heterozygous missense variant in the gene encoding E74 Like ETS Transcription Factor 2 (*ELF2*), which segregated with the disorder in three individuals in a single family.^47^ It is now apparent that the majority of individuals with CANVAS result from the homozygous inheritance of an expanded intronic pentamer in *RFC1*. We found the (AAGGG)_exp_ in 30 of 31 individuals with a RE at this locus. In only a single individual did we observe a different, presumably pathogenic motif; CANVAS8 had one allele with the (AAGGG)_exp_, whereas the second allele appeared to consist of an (AAAGG)_exp_. Notably, this alternate motif does not share the AAGGG haplotype (Figure S3). Analysis of the core haplotype in the majority of individuals with CANVAS suggests that the (AAGGG)_exp_ RE arose once, approximately 25,000 years ago, most likely in Europe. While the majority of individuals in our cohort who carry the (AAGGG)_n_ RE are of European ancestry, the RE is also present in non-European individuals, including a Lebanese family and two carriers of admixed Native American ancestry. Given the age of the CANVAS RE and recent human admixture it is likely that the locus may underlie CANVAS in apparently non-European individuals, despite the disorder being highly overrepresented in European populations.

Importantly, Cortese *et al* extend the clinical significance of the CANVAS RE by demonstrating it is potentially a common cause of unsolved ataxia not meeting the diagnostic criteria of CANVAS. Screening for homozygous inheritance of the (AAGGG)_exp_ RE in a cohort of 150 individuals with sporadic late-onset ataxia diagnosed 33 individuals (22%). This is consistent with the relatively high allele frequency of the (AAGGG)_exp_ we report in this paper. Collectively, the two studies screened for the RE in a total of 537 clinically normal samples, identifying 23 heterozygous and a single homozygous individual (allele frequency 25/1074=0.023). Given that the allele size and RE composition could not be determined in all controls, it is possible that the unaffected homozygous individual we identified carries two alleles smaller than the pathogenic range of >400 repeats reported by Cortese *et al*. However, the individual is less than half the mean age of CANVAS onset (∼60 years) and the lack of phenotype suggests the clinical features are yet to manifest.

### Mechanism of pathogenicity

There are multiple mechanisms by which RE can lead to pathogenicity, including RNA toxicity, protein toxicity and loss or gain of function.^7^ It is not yet known how the (AAGGG)_exp_ RE in *RFC1* causes CANVAS, however the homozygous inheritance pattern suggests a loss-of-function mechanism, rather than RNA or protein toxicity. In heterozygous carriers of the (AAGGG)_exp_ RE, our analysis of the GTEx RNA-seq data using haplotype tagging SNPs suggested that the pathogenic (AAGGG)_exp_ allele did not inhibit expression of *RFC1* compared to the reference (AAAAG)_11_ allele. Interestingly, Cortese *et al* also were unable to determine a mechanism of action. The (AAGGG)_exp_ RE did not appear to alter expression levels of *RFC1* or surrounding genes as determined by bulk RNAseq and qRT-PCR. Similarly, *RFC1* expression and protein levels appeared unchanged in peripheral or brain tissue derived from individuals with CANVAS, and no AAGGG RNA foci deposits were observed.^32^ While *RFC1* has not been previously associated with any disorder, it appears extremely intolerant to LoF (pLI = 0.97; observed/expected = 0.18, CI 0.12-0.31).^48^ In addition, siblings in the families studied carried the pathogenic RE in a heterozygous state but did not manifest any signs of the disorder. This observation is analogous to Friedreich’s ataxia, a recessive genetic ataxia caused by loss of function (LoF) of FRDA due to a pathogenic intronic RE.

*RFC1* encodes a subunit of replication factor C, a five-subunit protein complex required for DNA replication and repair. Analysis of the Genotype-Tissue Expression (GTEx) database demonstrated significant expression of *RFC1* in brain tissue, particularly the cerebellum. Replication factor C catalyzes opening the protein ring of proliferating cell nuclear antigen (PCNA), allowing it to encircle the DNA and function as a scaffold to recruit proteins involved in DNA replication, repair and remodeling.^49^ Mutations in multiple DNA replication and repair genes such as *TCD1, PNKP, XRCC1* and *APTX* result in ataxia^50^, highlighting the central role of this pathway in these overlapping disorders. One of the best known examples is the severe and early onset autosomal recessive disorder, ataxia telangiectasia, which is caused by mutations in the gene encoding ATM serine/threonine kinase (*ATM*), which is important for the repair of DNA double-strand breaks.^51^

The minimum pathogenic length and fine structure of the *RFC1* RE is currently unclear. While Cortese et al reported a pathogenic range of ∼400-2000, the individual repeat composition and a more precise repeat length was not determined. The short-read NGS technologies utilized in this study were unable to extend more than ∼100bp into the repeat sequence and efforts to amplify across the region using long range PCR were unsuccessful. While the repeat-primed PCR assay indicates the presence of the (AAGGG)_exp_ motif, it does not extend beyond ∼250bp (50 repeat units). The application of long read sequencing technologies currently being developed for RE disorders will be required to accurately elucidate both the length of the pathogenic allele and the repeat composition. Both of these parameters provide important clinical information regarding onset, progression and pathogenicity in other genetic ataxias such as SCA1 and Friedreich ataxia.^52;53^ Additional studies will also be required to elucidate the nature of the *RFC1* STR in control individuals. Cortese *et al* demonstrated considerable variability is present in the size and composition of the STR, but details regarding the size and composition of both normal and pathogenic alleles are yet to be fully determined. We show that the (AAGGG)_exp_ RE occurs within the 3 prime end of the Alu element, AluSx3. Alu elements typically have A-rich tails and in the reference sequence the *RFC1* Alu has an A-rich tail containing an (AAAAG)_11_ STR. There is some evidence that motifs that follow the pattern A_n_G_m_, especially (AAAG)_n_ and (AAAGG)_n_, display strong base-stacking interactions and are more likely to expand through replication slippage.^31^ This suggests an inherent mitotic instability of A and G rich motifs, consistent with what we observe in CANVAS. Notably, a number of pathogenic RE located with Alu have previously been described, including SCA10, SCA31, SCA37 and Friedreich ataxia.^22;26; 54; 55^

### Genomic re-diagnosis in CANVAS

Four of twenty two families enrolled in this study with a clinical diagnosis of CANVAS did not harbor the RE or any other potentially pathogenic variants in the *RFC1* locus. CANVAS13 was re-diagnosed with SCA3 after the WGS data was analyzed using our computational pipeline for detecting known pathogenic REs. In addition to cerebellar ataxia, individuals with SCA3 not uncommonly manifests a somatosensory impairment^56;57^ and vestibular involvement may be variably present^57^, resulting in a phenotype indistinguishable from CANVAS.^43^ This molecular re-diagnosis highlights the power of modern STR detection techniques to diagnose RE ataxias. In addition, NGS data provides the opportunity to simultaneously identify non-RE mediated causes of ataxia. In CANVAS17 we identified biallelic variants in *SACS* as the likely cause of disease. While individuals with spastic ataxia of the Charlevoix-Saguenay type may present with the combination of cerebellar ataxia and a peripheral neuropathy^58;59^ as seen in CANVAS, to our knowledge vestibular involvement has not previously been described, and so this potentially constitutes a novel manifestation of the disease. In addition, a very plausible heterozygous variant was identified in *FAT2* in CANVAS19. While classified as a VUS using ACMG guidelines,^60^ the variant is only observed once in gnomAD and was predicted pathogenic by multiple *in silico* algorithms. Very recently, heterozygous point mutations affecting the last cadherin domain (p.Lys3586Asn) or the linker region (p.Arg3649Gln) of *FAT2* have been associated with SCA45, adding weight to classifying the variant (p.Val1457Ala) in the thirteenth cadherin domain, as likely pathogenic. While the published clinical phenotype and mutation spectrum in SCA45 is limited, in common with CANVAS, it is a late onset and slowly progressive cerebellar ataxia.^45^

### Strengths and limitations of current STR detection tools

In this study, we implemented multiple computational tools to identify and validate the presence of a novel (AAGGG)_exp_ RE in the majority of individuals with a clinical diagnosis of CANVAS. In particular, the use of EHdn, with its non-reference based RE discovery framework, was crucial in identifying a putative candidate, with the reference-based STR detection tools facilitating the follow up analysis. Although all tools gave highly variable estimated repeat sizes, which are likely to be significantly less than the actual repeat size, they provided consistent evidence that the (AAGGG) motif was expanded. This level of evidence is helpful before embarking on the potentially complex process of molecular validation. In our analysis, only a single tool (GangSTR) failed to detect the alternate (AAAGG)_exp_ RE. It is not clear why this was the case, although it could be related to the more complicated [(AAAAG)_6_-(AAAGG)_exp_-(AAAAG)_6_] repeat structure. Notably, EHdn was able to identify the (AAAGG)_exp_ RE in CANVAS8-8, however genome wide significance was not achieved and the motif was less highly ranked based on p-value than the initial discovery of AAGGG (Table S3). This is likely due to the fact that the RE was only present in one copy and power was reduced as there was only a single case and a small number of controls. EHdn appears most effective as a discovery tool when used with multiple cases or larger numbers of controls. The results of this study highlight the importance of utilizing multiple tools to provide redundancy in the data analysis pipeline. We have now updated the exSTRa package (see weblinks below) to include CANVAS and other recently described pathogenic RE, providing additional utility to the research community to rapidly identify these RE in their cohorts. An additional issue we encountered, which potentially limited all tools, was the poor sequencing quality in reads containing the (AAGGG) motif compared to other STRs.

In conclusion, in this study we show that a recessively inherited, ancient RE located in intron 2 of *RFC1* is the predominant cause of CANVAS. Recently developed RE discovery tools facilitated the identification and verification of this novel RE, in addition to identifying other genetic causes of disease in the cohort. Despite the RE being located in an intron, we demonstrate that previously generated WES data with low-coverage genome-wide off target reads were helpful in providing increased statistical confidence in RE identification. Therefore, reanalysis of previously generated WES datasets potentially offers a cost effective approach to facilitating identification of novel intronic RE in discovery projects. Finally, we anticipate that implementation of these tools into routine diagnostic pipelines has the potential to significantly increase the current diagnostic rates of 36% and 17%, recorded for clinical exome and targeted panel analyses of individuals with ataxia, respectively.^61;62^

## SUPPLEMENTAL DATA

The supplemental data contain 7 figures and 4 tables.

## CONFLICTS OF INTEREST

The authors declare no conflicts of interest.

### ACKNOWLEDGMENTS

We acknowledge access to the dataset EGA00001003562 from the European Genome-Phenome Archive. This work was supported by the Australian Government National Health and Medical Research Council (Program Grant 1054618 to MB), the NIH (NINDS grant R01NS082094 to BLF) and the Murdoch Children’s Research Institute. MB was supported by an NHMRC Senior Research Fellowship (1102971) and DPB was supported by a Wellcome Clinical Research Career Development Fellowship. Additional funding was provided by the Independent Research Institute Infrastructure Support Scheme and the Victorian State Government Operational Infrastructure Program.

## WEB RESOURCES

exSTRa: https://github.com/bahlolab/exSTRa

Genotype-Tissue Expression (GTEx) project: https://gtexportal.org/home/

Genome Aggregation Database (gnomAD): http://gnomad.broadinstitute.org/

Integrative Genomics Viewer (IGV): http://software.broadinstitute.org/software/igv/

Online Mendelian Inheritance in Man: http://www.omim.org/

UCSC Genome Bioinformatics database: https://genome.ucsc.edu/

<u>Varsome: https://varsome.com</u>

## ACCESSION NUMBERS

The ClinVar details for the *RFC1* variants reported in this paper are accessible via submission SUB5220746.

## REFERENCES

1. McMurray, C.T. (2010). Mechanisms of trinucleotide repeat instability during human development. Nat Rev Genet 11, 786-799.

2. Gymrek, M., Willems, T., Guilmatre, A., Zeng, H., Markus, B., Georgiev, S., Daly, M.J., Price, A.L., Pritchard, J.K., Sharp, A.J., et al. (2016). Abundant contribution of short tandem repeats to gene expression variation in humans. Nature genetics 48, 22-29.

3. Quilez, J., Guilmatre, A., Garg, P., Highnam, G., Gymrek, M., Erlich, Y., Joshi, R.S., Mittelman, D., and Sharp, A.J. (2016). Polymorphic tandem repeats within gene promoters act as modifiers of gene expression and DNA methylation in humans. Nucleic Acids Res 44, 3750-3762.

4. Benson, G. (1999). Tandem repeats finder: a program to analyze DNA sequences. Nucleic Acids Res 27, 573-580.

5. Subramanian, S., Madgula, V.M., George, R., Mishra, R.K., Pandit, M.W., Kumar, C.S., and Singh, L. (2003). Triplet repeats in human genome: distribution and their association with genes and other genomic regions. Bioinformatics 19, 549-552.

6. La Spada, A.R., and Taylor, J.P. (2010). Repeat expansion disease: progress and puzzles in disease pathogenesis. Nat Rev Genet 11, 247-258.

7. Hannan, A.J. (2018). Tandem repeats mediating genetic plasticity in health and disease. Nat Rev Genet 19, 286-298.

8. Tankard, R.M., Bennett, M.F., Degorski, P., Delatycki, M.B., Lockhart, P.J., and Bahlo, M. (2018). Detecting Expansions of Tandem Repeats in Cohorts Sequenced with Short-Read Sequencing Data. American journal of human genetics 103, 858-873.

9. Ruano, L., Melo, C., Silva, M.C., and Coutinho, P. (2014). The global epidemiology of hereditary ataxia and spastic paraplegia: a systematic review of prevalence studies. Neuroepidemiology 42, 174-183.

10. Bird, T.D. (2018 Update). Hereditary Ataxia Overview. In GeneReviews((R)), M.P. Adam, H.H. Ardinger, R.A. Pagon, S.E. Wallace, L.J.H. Bean, K. Stephens, and A. Amemiya, eds. (Seattle (WA).

11. Szmulewicz, D.J., Waterston, J.A., Halmagyi, G.M., Mossman, S., Chancellor, A.M., McLean, C.A., and Storey, E. (2011). Sensory neuropathy as part of the cerebellar ataxia neuropathy vestibular areflexia syndrome. Neurology 76, 1903-1910.

12. Szmulewicz, D.J., McLean, C.A., MacDougall, H.G., Roberts, L., Storey, E., and Halmagyi, G.M. (2014). CANVAS an update: clinical presentation, investigation and management. J Vestib Res 24, 465-474.

13. Harding, A.E. (1981). “Idiopathic”late onset cerebellar ataxia. A clinical and genetic study of 36 cases. J Neurol Sci 51, 259-271.

14. Szmulewicz, D.J. (2017). Combined Central and Peripheral Degenerative Vestibular Disorders: CANVAS, Idiopathic Cerebellar Ataxia with Bilateral Vestibulopathy (CABV) and Other Differential Diagnoses of the CABV Phenotype. Curr Otorhinolaryngol Rep 5, 167–174.

15. Cha, Y.H. (2012). Less common neuro-otologic disorders. Continuum (Minneap Minn) 18, 1142-1157.

16. Szmulewicz, D.J., Merchant, S.N., and Halmagyi, G.M. (2011). Cerebellar ataxia with neuropathy and bilateral vestibular areflexia syndrome: a histopathologic case report. Otol Neurotol 32, e63-65.

17. Szmulewicz, D.J., McLean, C.A., Rodriguez, M.L., Chancellor, A.M., Mossman, S., Lamont, D., Roberts, L., Storey, E., and Halmagyi, G.M. (2014). Dorsal root ganglionopathy is responsible for the sensory impairment in CANVAS. Neurology 82, 1410-1415.

18. Szmulewicz, D.J., Seiderer, L., Halmagyi, G.M., Storey, E., and Roberts, L. (2015). Neurophysiological evidence for generalized sensory neuronopathy in cerebellar ataxia with neuropathy and bilateral vestibular areflexia syndrome. Muscle Nerve 51, 600-603.

19. Szmulewicz, D.J., Waterston, J.A., MacDougall, H.G., Mossman, S., Chancellor, A.M., McLean, C.A., Merchant, S., Patrikios, P., Halmagyi, G.M., and Storey, E. (2011). Cerebellar ataxia, neuropathy, vestibular areflexia syndrome (CANVAS): a review of the clinical features and video-oculographic diagnosis. Ann N Y Acad Sci 1233, 139-147.

20. Petersen, J.A., Wichmann, W.W., and Weber, K.P. (2013). The pivotal sign of CANVAS. Neurology 81, 1642-1643.

21. Szmulewicz, D., MacDougall, H., Storey, E., Curthoys, I., and Halmagyi, M. (2014). A Novel Quantitative Bedside Test of Balance Function: The Video Visually Enhanced Vestibulo-ocular Reflex (VVOR) Neurology 82, S19.002.

22. Campuzano, V., Montermini, L., Molto, M.D., Pianese, L., Cossee, M., Cavalcanti, F., Monros, E., Rodius, F., Duclos, F., Monticelli, A., et al. (1996). Friedreich’s ataxia: autosomal recessive disease caused by an intronic GAA triplet repeat expansion. Science 271, 1423-1427.

23. Taki, M., Nakamura, T., Matsuura, H., Hasegawa, T., Sakaguchi, H., Morita, K., Ishii, R., Mizuta, I., Kasai, T., Mizuno, T., et al. (2018). Cerebellar ataxia with neuropathy and vestibular areflexia syndrome (CANVAS). Auris Nasus Larynx 45, 866-870.

24. Maruta, K., Aoki, M., and Sonoda, Y. (2019). [Cerebellar ataxia with neuropathy and vestibular areflexia syndrome (CANVAS): a case report]. Rinsho Shinkeigaku 59, 27-32.

25. Bahlo, M., Bennett, M.F., Degorski, P., Tankard, R.M., Delatycki, M.B., and Lockhart, P.J. (2018). Recent advances in the detection of repeat expansions with short-read next-generation sequencing. F1000Res 7.

26. Seixas, A.I., Loureiro, J.R., Costa, C., Ordonez-Ugalde, A., Marcelino, H., Oliveira, C.L., Loureiro, J.L., Dhingra, A., Brandao, E., Cruz, V.T., et al. (2017). A Pentanucleotide ATTTC Repeat Insertion in the Non-coding Region of DAB1, Mapping to SCA37, Causes Spinocerebellar Ataxia. American journal of human genetics 101, 87-103.

27. Ishiura, H., Doi, K., Mitsui, J., Yoshimura, J., Matsukawa, M.K., Fujiyama, A., Toyoshima, Y., Kakita, A., Takahashi, H., Suzuki, Y., et al. (2018). Expansions of intronic TTTCA and TTTTA repeats in benign adult familial myoclonic epilepsy. Nature genetics 50, 581-590.

28. Dolzhenko, E., van Vugt, J., Shaw, R.J., Bekritsky, M.A., van Blitterswijk, M., Narzisi, G., Ajay, S.S., Rajan, V., Lajoie, B.R., Johnson, N.H., et al. (2017). Detection of long repeat expansions from PCRfree whole-genome sequence data. Genome research 27, 1895-1903.

29. Tang, H., Kirkness, E.F., Lippert, C., Biggs, W.H., Fabani, M., Guzman, E., Ramakrishnan, S., Lavrenko, V., Kakaradov, B., Hou, C., et al. (2017). Profiling of Short-Tandem-Repeat Disease Alleles in 12,632 Human Whole Genomes. American journal of human genetics 101, 700-715.

30. Dashnow, H., Lek, M., Phipson, B., Halman, A., Sadedin, S., Lonsdale, A., Davis, M., Lamont, P., Clayton, J.S., Laing, N.G., et al. (2018). STRetch: detecting and discovering pathogenic short tandem repeat expansions. Genome biology 19, 121.

31. Mousavi, N., Shleizer-Burko, S., and Gymrek, M. (2018). Profiling the genome-wide landscape of tandem repeat expansions. BioRxiv, https://doi.org/10.1101/361162

32. Cortese, A., Simone, R., Sullivan, R., Vandrovcova, J., Tariq, H., Yan, Y.W., Humphrey, J., Jaunmuktane, Z., Sivakumar, P., Polke, J., et al. (2019). Biallelic expansion of an intronic repeat in RFC1 is a common cause of late-onset ataxia. Nature genetics 51, 649-658.

33. Smith, K.R., Bromhead, C.J., Hildebrand, M.S., Shearer, A.E., Lockhart, P.J., Najmabadi, H., Leventer, R.J., McGillivray, G., Amor, D.J., Smith, R.J., et al. (2011). Reducing the exome search space for mendelian diseases using genetic linkage analysis of exome genotypes. Genome biology 12, R85.

34. Bahlo, M., and Bromhead, C.J. (2009). Generating linkage mapping files from Affymetrix SNP chip data. Bioinformatics 25, 1961-1962.

35. Abecasis, G.R., Cherny, S.S., Cookson, W.O., and Cardon, L.R. (2002). Merlin-rapid analysis of dense genetic maps using sparse gene flow trees. Nature genetics 30, 97-101.

36. Quinlan, A.R., and Hall, I.M. (2010). BEDTools: a flexible suite of utilities for comparing genomic features. Bioinformatics 26, 841-842.

37. Pedersen, B.S., Layer, R.M., and Quinlan, A.R. (2016). Vcfanno: fast, flexible annotation of genetic variants. Genome biology 17, 118.

38. Wang, K., Li, M., and Hakonarson, H. (2010). ANNOVAR: functional annotation of genetic variants from high-throughput sequencing data. Nucleic Acids Res 38, e164.

39. Dobin, A., Davis, C.A., Schlesinger, F., Drenkow, J., Zaleski, C., Jha, S., Batut, P., Chaisson, M., and Gingeras, T.R. (2013). STAR: ultrafast universal RNA-seq aligner. Bioinformatics 29, 15-21.

40. Liao, Y., Smyth, G.K., and Shi, W. (2014). featureCounts: an efficient general purpose program for assigning sequence reads to genomic features. Bioinformatics 30, 923-930.

41. Ritchie, M.E., Phipson, B., Wu, D., Hu, Y., Law, C.W., Shi, W., and Smyth, G.K. (2015). limma powers differential expression analyses for RNA-sequencing and microarray studies. Nucleic Acids Res 43, e47.

42. Gandolfo, L.C., Bahlo, M., and Speed, T.P. (2014). Dating rare mutations from small samples with dense marker data. Genetics 197, 1315-1327.

43. Szmulewicz, D.J., Roberts, L., McLean, C.A., MacDougall, H.G., Halmagyi, G.M., and Storey, E. (2016). Proposed diagnostic criteria for cerebellar ataxia with neuropathy and vestibular areflexia syndrome (CANVAS). Neurol Clin Pract 6, 61-68.

44. Wang, K., Li, M., Hadley, D., Liu, R., Glessner, J., Grant, S.F., Hakonarson, H., and Bucan, M. (2007). PennCNV: an integrated hidden Markov model designed for high-resolution copy number variation detection in whole-genome SNP genotyping data. Genome research 17, 1665-1674.

45. Nibbeling, E.A.R., Duarri, A., Verschuuren-Bemelmans, C.C., Fokkens, M.R., Karjalainen, J.M., Smeets, C., de Boer-Bergsma, J.J., van der Vries, G., Dooijes, D., Bampi, G.B., et al. (2017). Exome sequencing and network analysis identifies shared mechanisms underlying spinocerebellar ataxia. Brain 140, 2860&2878.

46. Rinne, T., Bronstein, A.M., Rudge, P., Gresty, M.A., and Luxon, L.M. (1998). Bilateral loss of vestibular function: clinical findings in 53 patients. J Neurol 245, 314&321.

47. Ahmad, H., Requena, T., Frejo, L., Cobo, M., Gallego-Martinez, A., Martin, F., Lopez-Escamez, J.A., and Bronstein, A.M. (2018). Clinical and Functional Characterization of a Missense ELF2 Variant in a CANVAS Family. Front Genet 9, 85.

48. Lek, M., Karczewski, K.J., Minikel, E.V., Samocha, K.E., Banks, E., Fennell, T., O’Donnell-Luria, A.H., Ware, J.S., Hill, A.J., Cummings, B.B., et al. (2016). Analysis of protein-coding genetic variation in 60,706 humans. Nature 536, 285&291.

49. Zhang, G., Gibbs, E., Kelman, Z., O’Donnell, M., and Hurwitz, J. (1999). Studies on the interactions between human replication factor C and human proliferating cell nuclear antigen. Proc Natl Acad Sci U S A 96, 1869&1874.

50. Yoon, G., and Caldecott, K.W. (2018). Nonsyndromic cerebellar ataxias associated with disorders of DNA single-strand break repair. Handbook of clinical neurology 155, 105&115.

51. Savitsky, K., Bar-Shira, A., Gilad, S., Rotman, G., Ziv, Y., Vanagaite, L., Tagle, D.A., Smith, S., Uziel, T., Sfez, S., et al. (1995). A single ataxia telangiectasia gene with a product similar to PI-3 kinase. Science 268, 1749&1753.

52. Kraus-Perrotta, C., and Lagalwar, S. (2016). Expansion, mosaicism and interruption: mechanisms of the CAG repeat mutation in spinocerebellar ataxia type 1. Cerebellum Ataxias 3, 20.

53. Mateo, I., Llorca, J., Volpini, V., Corral, J., Berciano, J., and Combarros, O. (2003). GAA expansion size and age at onset of Friedreich’s ataxia. Neurology 61, 274&275.

54. Bushara, K., Bower, M., Liu, J., McFarland, K.N., Landrian, I., Hutter, D., Teive, H.A., Rasmussen, A., Mulligan, C.J., and Ashizawa, T. (2013). Expansion of the Spinocerebellar ataxia type 10 (SCA10) repeat in a patient with Sioux Native American ancestry. PloS one 8, e81342.

55. Sato, N., Amino, T., Kobayashi, K., Asakawa, S., Ishiguro, T., Tsunemi, T., Takahashi, M., Matsuura, T., Flanigan, K.M., Iwasaki, S., et al. (2009). Spinocerebellar ataxia type 31 is associated with “inserted”penta-nucleotide repeats containing (TGGAA)n. American journal of human genetics 85, 544&557.

56. Jardim, L.B., Pereira, M.L., Silveira, I., Ferro, A., Sequeiros, J., and Giugliani, R. (2001). Neurologic findings in Machado-Joseph disease: relation with disease duration, subtypes, and (CAG)n. Arch Neurol 58, 899&904.

57. Gordon, C.R., Zivotofsky, A.Z., and Caspi, A. (2014). Impaired vestibulo-ocular reflex (VOR) in spinocerebellar ataxia type 3 (SCA3): bedside and search coil evaluation. J Vestib Res 24, 351&355.

58. Gagnon, C., Desrosiers, J., and Mathieu, J. (2004). Autosomal recessive spastic ataxia of Charlevoix-Saguenay: upper extremity aptitudes, functional independence and social participation. Int J Rehabil Res 27, 253&256.

59. Vill, K., Muller-Felber, W., Glaser, D., Kuhn, M., Teusch, V., Schreiber, H., Weis, J., Klepper, J., Schirmacher, A., Blaschek, A., et al. (2018). SACS variants are a relevant cause of autosomal recessive hereditary motor and sensory neuropathy. Human genetics 137, 911&919.

60. Richards, S., Aziz, N., Bale, S., Bick, D., Das, S., Gastier-Foster, J., Grody, W.W., Hegde, M., Lyon, E., Spector, E., et al. (2015). Standards and guidelines for the interpretation of sequence variants: a joint consensus recommendation of the American College of Medical Genetics and Genomics and the Association for Molecular Pathology. Genet Med 17, 405&424.

61. Sullivan, R., Yau, W.Y., O’Connor, E., and Houlden, H. (2019). Spinocerebellar ataxia: an update. J Neurol 266, 533&544.

62. Galatolo, D., Tessa, A., Filla, A., and Santorelli, F.M. (2018). Clinical application of next generation sequencing in hereditary spinocerebellar ataxia: increasing the diagnostic yield and broadening the ataxia-spasticity spectrum. A retrospective analysis. Neurogenetics 19, 1&8.

