## Supplemental for "Validation of new bioinformatic tools to identify expanded repeats: a non-reference intronic pentamer expansion in *RFC1* causes CANVAS"

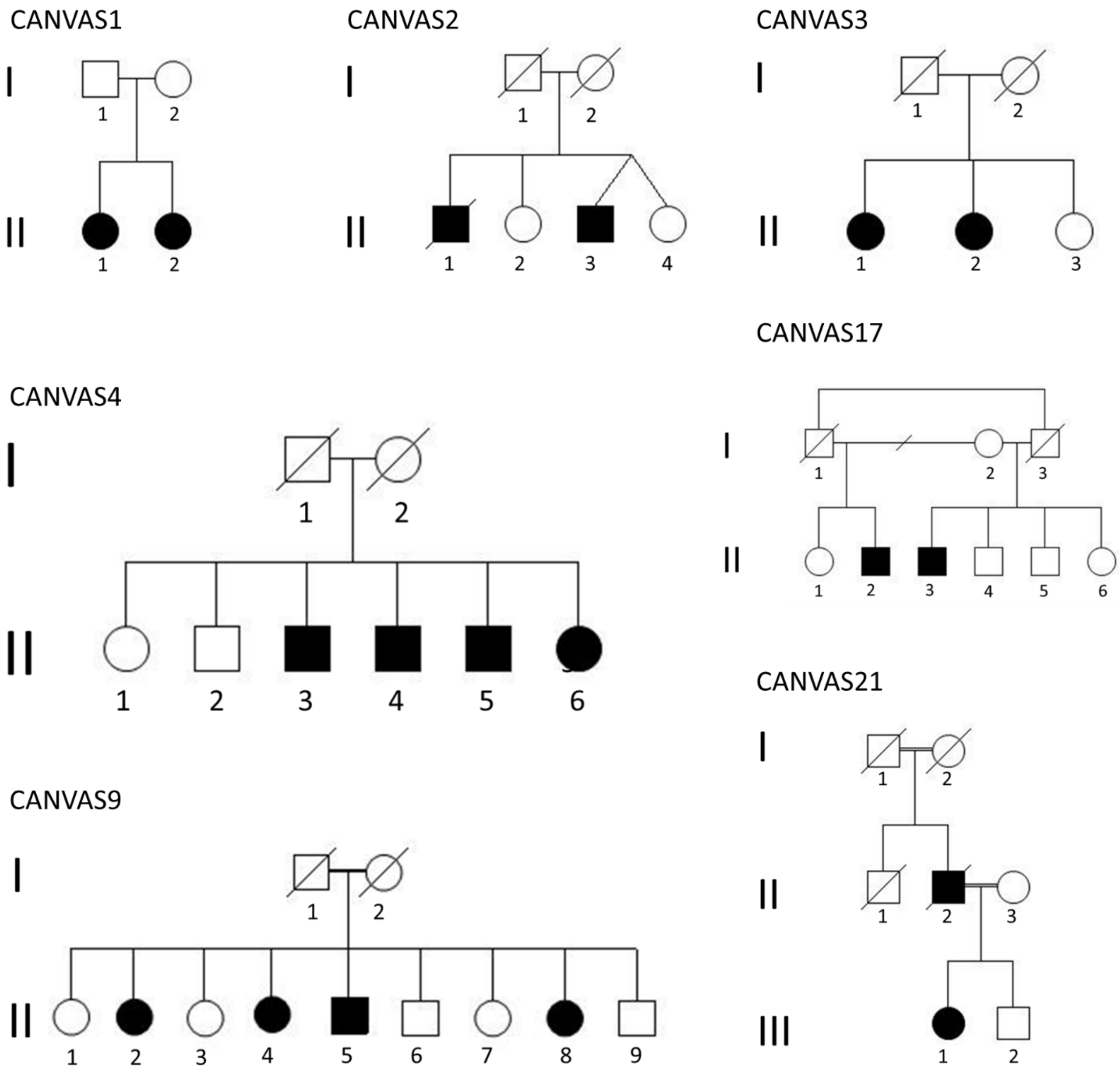


**Figure S1: Pedigree structure of CANVAS families utilized for linkage studies (CANVAS1, 2, 3, 4 and 9) or demonstrating multigenerational inheritance.**

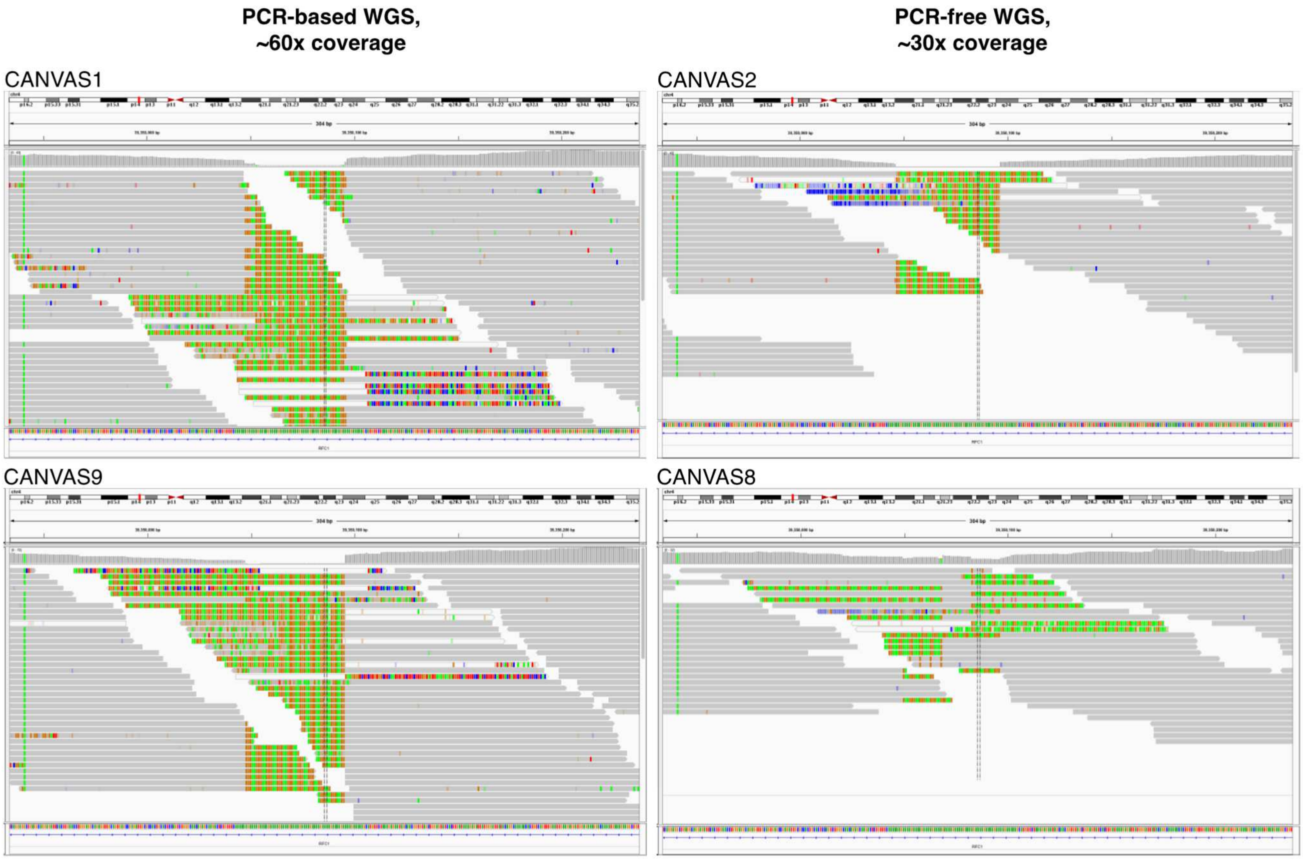


**Figure S2: IGV snapshots of the (AAGGG)n locus in *RFC1*.**

Illustrated are affected individuals from homozygous carriers of AAGGG (CANVAS1, 2, 9) and an individual who carries the AAGGG allele in addition to an expanded AAAGG allele (CANVAS8).

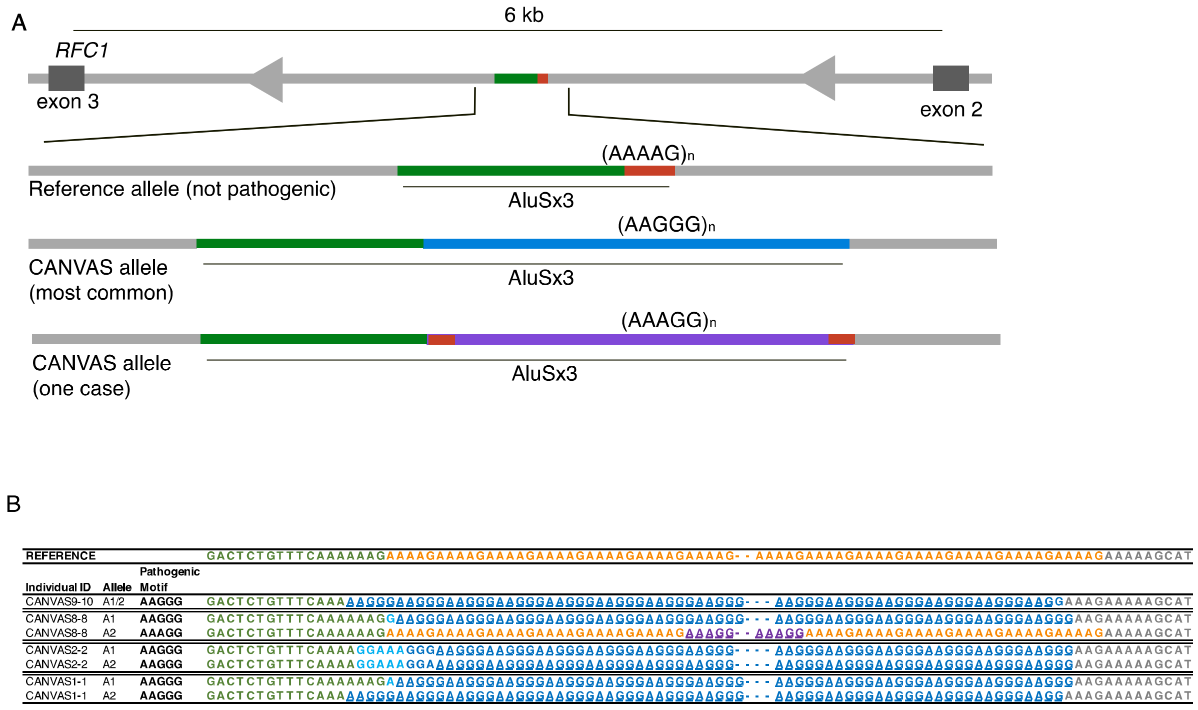


**Figure S3: Schematic representation of pathogenic CANVAS alleles.**

(A) Visual representation of the AAGGG and AAAGG repeat expansions in the second intron of *RFC1*. (B) Visual representation of the different AAGGG/AAAGG alleles in affected individuals identified by WGS.

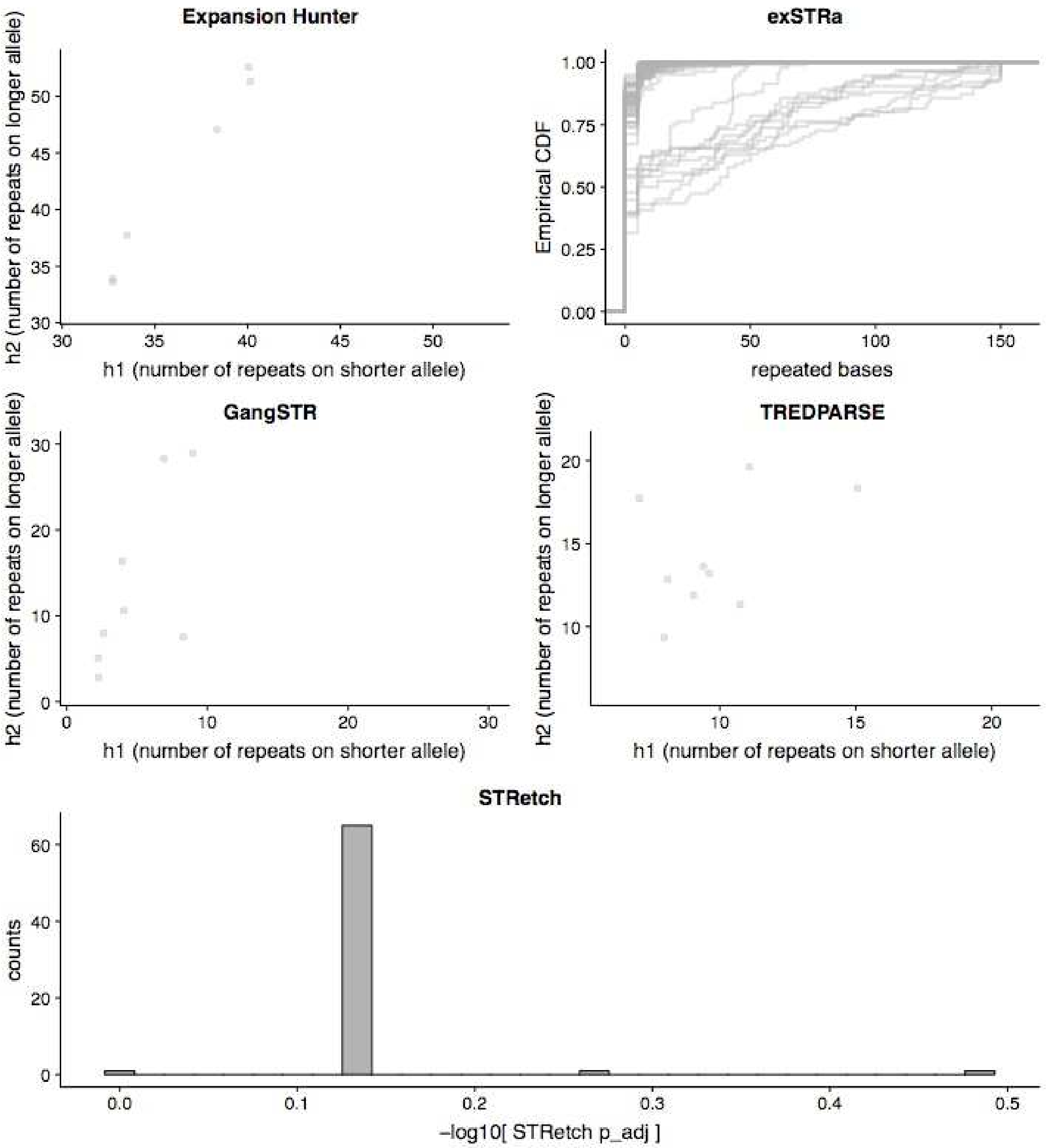


**Figure S4: Computational validation of the $(AAGGG)_n$ STR.**

The WGS from 69 unrelated non-CANVAS individuals (Coriell dataset) were analysed at the coordinate's chr4:39350045-39350095, using the tools exSTRa, EH, GangSTR, TREDPARSE and STRetch.

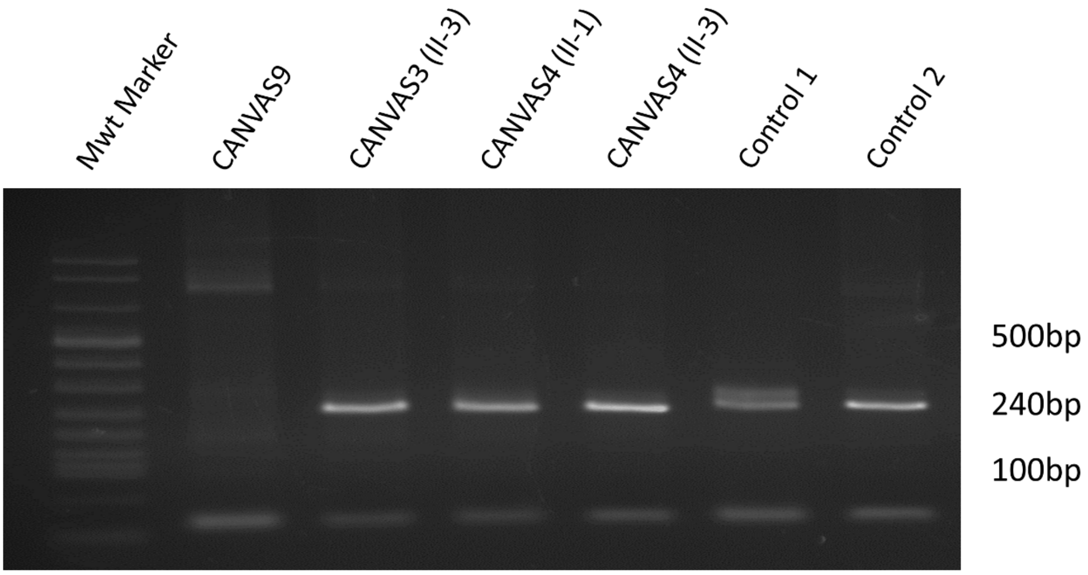


**Figure S5: PCR analysis of unaffected individuals from *RFC1* linked CANVAS families.** Genomic DNA from three unaffected family members from CANVAS3 and CANVAS4 was analysed for the presence of a non-expanded *RFC1* allele. All three individuals carried at least one non-expanded allele, as indicated by the ~250bp PCR product. CANVAS9 is homozygous for the expanded allele and indicates the pattern expected in the absence of the reference allele. Pedigree structure and affected status are illustrated in Figure S1.

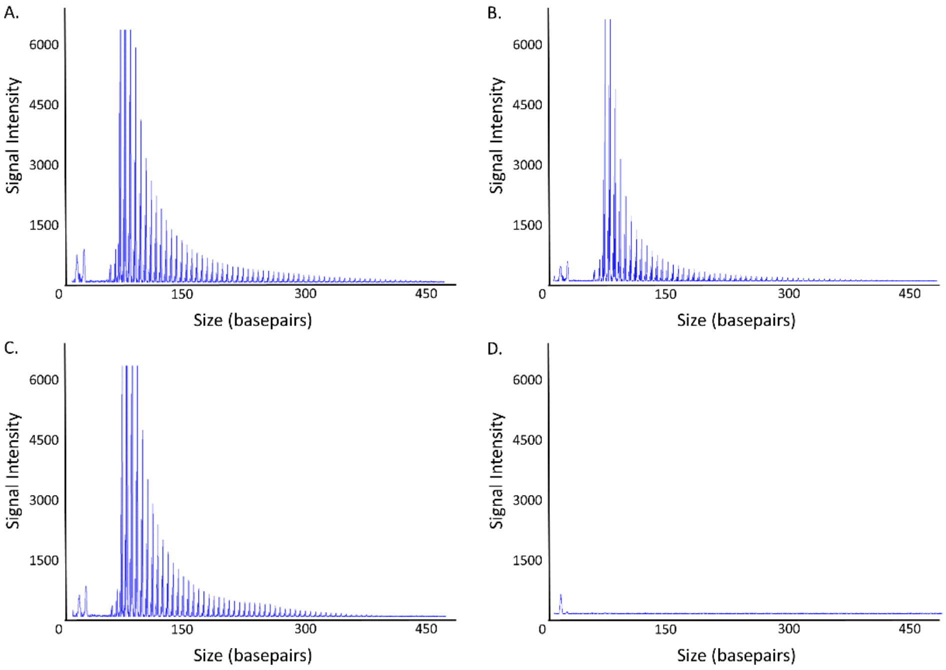


**Figure S6: Repeat-primed PCR analysis of unaffected individuals with (AAGGG)exp RE.** Representative images of the repeat-primed PCR for the (AAGGG)exp RE demonstrating a saw-toothed product with 5 base pair repeat unit size, amplified from gDNA of control individuals with heterozygous (A, B) or homozygous (C) alleles encoding the (AAGGG)exp RE. No product was observed for the no gDNA template negative control (D).

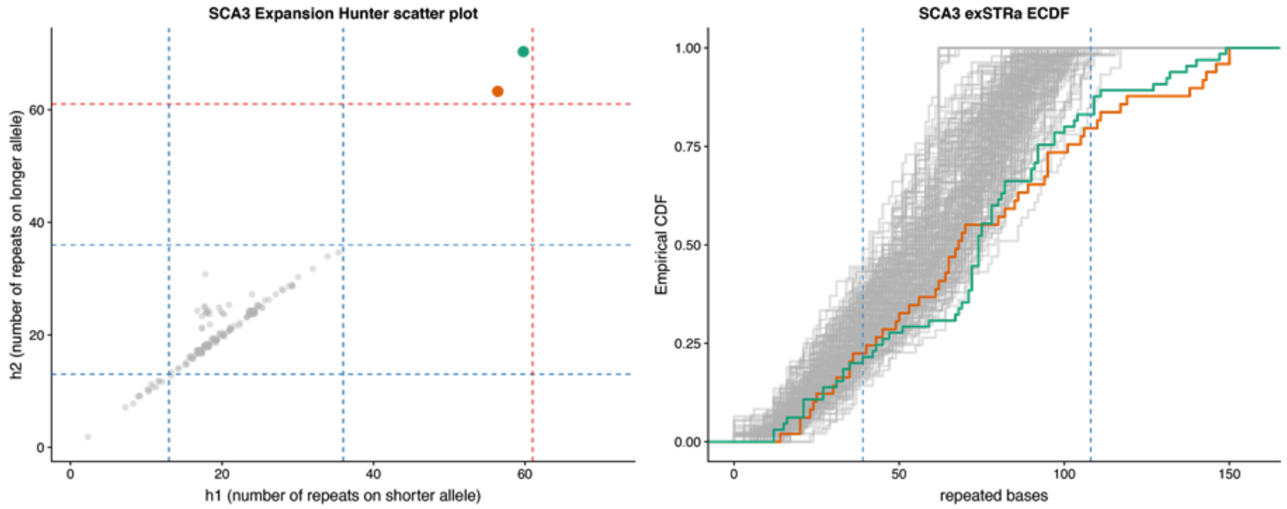


**Figure S7: Identification of a pathogenic SCA3 expansion in CANVAS13.**

The WGS data for SCA3 (green dot) and an individual with a confirmed SCA3 RE (orange dot) was analysed using ExpansionHunter and exSTRa. This analysis identified a heterozygous, pathogenic SCA3 expansion in CANVAS13.

| Primer name | Position (hg19) | Sequence |
|---|---|---|
| CANVAS_RFC1_3F | chr4: 39350172-39350192 | 5'-ACTGACAGTGTTTTTGCCTGT |
| CANVAS_RFC1_3R | chr4:39349940-39349959 | 5'-GGCTGAGGCAGGAGATTCAC |
| TPP_CANVAS_FAM_2F | chr4:39350172-39350192 | FAM-5'-ACTGACAGTGTTTTTGCCTGT |
| 5R_TPP_M13R_CANVAS_RE_R | NA | 5'-CAGGAAACAGCTATGACC_AAGGGAAGGGAAGGGAAGGGAAGGG |
| TPP_M13R | NA | 5'-CAGGAAACAGCTATGACC |

**Table S1: Primer sequences for analysis of RFC1.**

The gene reference sequences utilized were NC_000004 and NM_002913 (*RFC1*).

| ENSEMBL Gene ID | Gene name | Gene biotype | OMIM gene ID | OMIM disease phenotype (phenotype, OMIM phenotype ID, inheritance pattern) |
|---|---|---|---|---|
| ENSG00000121895 | TMEM156 | protein_coding | NA | NA |
| ENSG00000109790 | KLHL5 | protein_coding | 608064 | NA |
| ENSG00000249207 | RP11-360F5.1 | antisense | NA | NA |
| ENSG00000249685 | RP11-360F5.3 | lincRNA | NA | NA |
| ENSG00000157796 | WDR19 | protein_coding | 608151 | ?Cranioectodermal dysplasia 4, 614378, AR; ?Short-rib thoracic dysplasia 5 with or without polydactyly, 614376, AR;Nephronophthisis 13, 614377, AR; Senior-Loken syndrome 8, 616307, AR |
| ENSG00000035928 | RFC1 | protein_coding | 102579 | NA |
| ENSG00000206675 | RNU6-32P | snRNA | NA | NA |
| ENSG00000222592 | RNU6-887P | snRNA | NA | NA |
| ENSG00000134962 | KLB | protein_coding | 611135 | NA |
| ENSG00000264621 | MIR5591 | miRNA | NA | NA |
| ENSG00000238797 | Y_RNA | misc_RNA | NA | NA |
| ENSG00000163682 | RPL9 | protein_coding | 603686 | NA |
| ENSG00000121897 | LIAS | protein_coding | 607031 | Hyperglycinemia, lactic acidosis, and seizures, 614462, AR |
| ENSG00000224097 | RP11-472B18.1 | pseudogene | NA | NA |
| ENSG00000109814 | UGDH | protein_coding | 603370 | NA |
| ENSG00000249348 | UGDH-AS1 | antisense | NA | NA |
| ENSG00000163683 | SMIM14 | protein_coding | NA | NA |
| ENSG00000252796 | RNU7-11P | snRNA | NA | NA |
| ENSG00000255458 | RP11-539G18.2 | lincRNA | NA | NA |
| ENSG00000078140 | UBE2K | protein_coding | 602846 | NA |
| ENSG00000252975 | Y_RNA | misc_RNA | NA | NA |
| ENSG00000249019 | RP11-539G18.1 | pseudogene | NA | NA |
| ENSG00000243260 | RN7SL558P | misc_RNA | NA | NA |
| ENSG00000180610 | ZBTB12P1 | pseudogene | NA | NA |

| | | | | |
|---|---|---|---|---|
| ENSG00000121892 | PDS5A | protein_coding | 613200 | NA |
| ENSG00000271278 | TCEB1P33 | pseudogene | NA | NA |
| ENSG00000252970 | RNA5SP159 | rRNA | NA | NA |
| ENSG00000250568 | RP11-333E13.2 | pseudogene | NA | NA |
| ENSG00000231707 | PABPC1P1 | pseudogene | NA | NA |
| ENSG00000249064 | KRT18P25 | pseudogene | NA | NA |
| ENSG00000205794 | RP11-333E13.4 | pseudogene | NA | NA |
| ENSG00000078177 | N4BP2 | protein_coding | NA | NA |
| ENSG00000200455 | RNU6-1112P | snRNA | NA | NA |
| ENSG00000201863 | SNORA51 | snoRNA | NA | NA |
| ENSG00000248977 | RP11-395I6.1 | pseudogene | NA | NA |
| ENSG00000260296 | RP11-395I6.3 | sense_overlapping | NA | NA |
| ENSG00000168421 | RHOH | protein_coding | 602037 | {?Epidermodysplasia verruciformis, susceptibility to, 4}, 618307, AR |
| ENSG00000250338 | RP11-395I6.2 | lincRNA | NA | NA |
| ENSG00000249241 | AC195454.1 | lincRNA | NA | NA |
| ENSG00000174343 | CHRNA9 | protein_coding | 605116 | NA |
| ENSG00000239010 | RNU7-74P | snRNA | NA | NA |
| ENSG00000250893 | RP11-588L15.2 | antisense | NA | NA |

**Table S2. Genes and known disease associations within the CANVAS linkage region**

| region | repeat. | pval | adj | Func.refGene | Gene.refGene |
|---|---|---|---|---|---|
| chr4:39350095-39350508 | AAGGG | 0.0027 | 1 | intronic | RFC1 |
| chr5:68499727-68500146 | AATATATATATATAG | 0.0027 | 1 | intronic | CENPH |
| chr7:157344383-157344946 | ACAGCCACCACCCACCCC | 0.0027 | 1 | intronic | PTPRN2 |
| chr9:138739792-138740277 | AAGGGGAGGGGAGTGGGGGG | 0.0027 | 1 | intronic | CAMSAP1 |
| chr11:100495523-100496852 | AATATGTGTATATATGT | 0.0027 | 1 | intergenic | CNTN5; LOC100128386 |
| chr11:127149089-127150146 | AAG | 0.0027 | 1 | ncRNA_intronic | LOC101929497 |
| chr19:43901669-43901885 | AAATATATATTATATAT | 0.0027 | 1 | intronic | TEX101 |
| chr20:43066951-43067388 | AAAAATATAATATAT | 0.0027 | 1 | intergenic | HNF4A; LINC01430 |
| chr3:175416044-175416658 | ACACATACATATAT | 0.0028 | 1 | intronic | NAALADL2 |
| chr5:4286195-4286524 | AAGCTATATATATATATAGTG | 0.0028 | 1 | intergenic | IRX1; LINC02114 |
| chr5:74613999-74614513 | AAAG | 0.0028 | 1 | intergenic | ANKRD31; HMGCR |
| chr6:165352545-165353525 | AAAG | 0.0028 | 1 | intergenic | MEAT6; C6orf118 |
| chr7:4930864-4931703 | AGAT | 0.0028 | 1 | intergenic | RADIL; MMD2 |
| chr7:157846063-157847123 | ACCCCAGAGACGCAGAG | 0.0028 | 1 | intronic | PTPRN2 |
| chr10:134789654-134790481 | AATACATTCCACGTGTATC | 0.0028 | 1 | ncRNA_exonic | LINC01168 |
| chr11:2182294-2182840 | ACACCCCTGTCCCC | 0.0028 | 1 | UTR5 | INS; INS-IGF2 |
| chr11:120746805-120747444 | ACC | 0.0028 | 1 | ncRNA_intronic | LOC101929227 |
| chr11:134177695-134179032 | ACC | 0.0028 | 1 | intronic | GLB1L3 |
| chr13:40788773-40788913 | AATATAT | 0.0028 | 1 | ncRNA_intronic | LINC00548 |

**Table S3: Expansion Hunter de novo results - screening for RE in two CANVAS individuals compared to 31 controls (all PCR based WGS, $p < 0.01$) ranked by p-value and ordered by chromosome**

| position | REF | ALT | genes | region | SNP | MAF | CANVAS1 | CANVAS2 | CANVAS3 | CANVAS4 | CANVAS6 | CANVAS7 | CANVAS8 | CANVAS9 | CANVAS10 | CANVAS12 | CANVAS18 | CANVAS14 | Core haplotype |
|---|---|---|---|---|---|---|---|---|---|---|---|---|---|---|---|---|---|---|---|
| 38777173 | A | T | TLR10 | exonic | rs10856838 | 0.2806 | 0/0 | | 0/0 | 0/1 | 0/1 | 1/1 | 0/0 | 0/0 | 0/0 | 0/1 | 0/1 | 0/0 | |
| 38777236 | T | G | TLR10 | UTR5 | rs10856839 | 0.2803 | 0/0 | 0/0 | 0/0 | 0/1 | 0/1 | 1/1 | 0/0 | 0/0 | 0/0 | 0/1 | 0/1 | 0/0 | |
| 38798515 | G | A | TLR1 | exonic | rs72493538 | 0.0063 | 0/0 | 0/0 | 0/0 | 0/0 | 0/0 | 0/1 | 0/0 | 0/0 | 0/0 | 0/0 | 0/0 | 0/0 | |
| 38798648 | C | A | TLR1 | exonic | rs5743618 | 0.4732 | 0/0 | 0/1 | 0/0 | 1/1 | 0/0 | 1/1 | 0/0 | 1/1 | 0/0 | 1/1 | 1/1 | 1/1 | |
| 38798935 | C | T | TLR1 | exonic | rs5743614 | 0.3942 | 0/0 | 0/1 | 0/0 | 1/1 | 0/0 | 0/1 | 0/0 | 0/0 | 0/0 | 1/1 | 0/1 | 1/1 | |
| 38799269 | G | A | TLR1 | exonic | rs770320905 | 3.23E-05 | 0/0 | 0/0 | 0/0 | 0/1 | 0/0 | 0/0 | 0/0 | 0/0 | 0/0 | 0/0 | 0/0 | 0/0 | |
| 38799539 | T | A | TLR1 | exonic | rs3923647 | 0.0364 | 0/0 | 0/0 | 0/0 | 0/0 | 0/0 | 0/0 | 0/0 | 0/0 | 0/0 | 0/0 | 0/0 | 1/1 | |
| 38799710 | T | C | TLR1 | exonic | rs4833095 | 0.4041 | 0/0 | 0/1 | 0/0 | 1/1 | 0/0 | 0/1 | 0/0 | 0/0 | 0/0 | 1/1 | 0/1 | 1/1 | |
| 38800214 | C | G | TLR1 | exonic | rs5743611 | 0.1029 | 0/0 | 0/0 | 0/0 | 0/0 | 0/1 | 0/0 | 0/0 | 0/0 | 0/0 | 0/0 | 0/0 | 0/0 | |
| 38829163 | A | C | TLR6 | exonic | rs5743818 | 0.2286 | 1/1 | 0/0 | 0/1 | 0/0 | 0/1 | 0/0 | 0/0 | 0/0 | 0/0 | 0/0 | 0/0 | 0/0 | |
| 38829832 | T | C | TLR6 | exonic | rs3775073 | 0.4618 | 1/1 | 0/0 | 0/1 | 0/1 | 0/1 | 0/0 | 0/0 | 0/0 | 0/0 | 0/1 | 0/1 | 1/1 | |
| 38830012 | G | C | TLR6 | exonic | rs3821985 | 0.4546 | 1/1 | 0/0 | 0/1 | 0/1 | 0/1 | 0/0 | 0/0 | 0/0 | 0/0 | 0/1 | 0/1 | 1/1 | |
| 38830116 | C | T | TLR6 | exonic | rs3796508 | 0.0134 | 0/0 | 0/0 | 0/0 | 0/1 | 0/0 | 0/0 | 0/0 | 0/0 | 0/0 | 0/1 | 0/1 | 0/0 | |
| 38830350 | A | G | TLR6 | exonic | rs5743810 | 0.7116 | 1/1 | 0/1 | 1/1 | 1/1 | 0/1 | 1/1 | 0/0 | 0/0 | 0/0 | 1/1 | 1/1 | 1/1 | |
| 38830736 | A | G | TLR6 | exonic | rs5743808 | 0.0336 | 0/0 | 0/0 | 0/0 | 0/1 | 0/0 | 0/0 | 0/0 | 0/0 | 0/0 | 0/1 | 0/1 | 0/0 | |
| 38879655 | C | T | FAM114A1 | intronic | rs73236661 | 0.3071 | 1/1 | 0/0 | 0/1 | 0/1 | 0/1 | 0/0 | 0/0 | 0/0 | 0/0 | 0/0 | 0/1 | 0/0 | |
| 38879949 | G | A | FAM114A1 | exonic | rs11096964 | 0.308 | 1/1 | 0/0 | 0/1 | 0/1 | 0/1 | 0/0 | 0/0 | 0/0 | 0/0 | 0/0 | 0/1 | 0/0 | |
| 38880046 | T | C | FAM114A1 | exonic | rs11555334 | 0.3075 | 1/1 | 0/0 | 0/1 | 0/1 | 0/1 | 0/0 | 0/0 | 0/0 | 0/0 | 0/0 | 0/1 | 0/0 | |
| 38880136 | A | T | FAM114A1 | intronic | rs11943209 | 0.308 | 1/1 | 0/0 | 0/1 | 0/1 | 0/1 | 0/0 | 0/0 | 0/0 | 0/0 | 0/0 | 0/1 | 0/0 | |
| 38880167 | C | T | FAM114A1 | intronic | rs11938589 | 0.308 | 1/1 | 0/0 | 0/1 | 0/1 | 0/1 | 0/0 | 0/0 | 0/0 | 0/0 | 0/0 | 0/1 | 0/0 | |
| 38880181 | A | ATGTGT | FAM114A1 | intronic | rs111676993 | 0.3053 | 1/1 | 0/0 | 0/1 | 0/1 | 0/1 | 0/0 | 0/0 | 0/0 | 0/0 | 0/0 | 0/1 | 0/0 | |
| 38924358 | A | G | FAM114A1 | intronic | rs12498685 | 0.1954 | 1/1 | 0/0 | 0/1 | 0/0 | 0/0 | 1/1 | 0/0 | 0/0 | 0/0 | 0/0 | 0/0 | 0/0 | |
| 38930921 | A | G | FAM114A1 | exonic | rs3188469 | 0.1131 | 1/1 | 0/0 | 0/1 | 0/0 | 0/0 | 0/0 | 0/0 | 0/0 | 0/0 | 0/0 | 0/0 | 0/0 | |
| 38930985 | T | TAA | FAM114A1 | intronic | rs35369040 | 0.2194 | 0/0 | 0/0 | 0/1 | 0/1 | 0/0 | 0/0 | 0/0 | 1/1 | 0/0 | 0/0 | 0/0 | 0/0 | |
| 38930995 | C | A | FAM114A1 | intronic | rs58035134 | 0.2307 | 0/0 | 0/0 | 0/1 | 0/1 | 0/0 | 0/0 | 0/0 | 1/1 | 0/0 | 0/0 | 0/0 | 0/0 | |
| 38937372 | T | C | FAM114A1 | exonic | rs2271031 | 0.6742 | 1/1 | 0/0 | 1/1 | 1/1 | 1/1 | 1/1 | 0/1 | 1/1 | 0/0 | 0/0 | 0/0 | 0/0 | |
| 38944976 | A | AT | FAM114A1 | intronic | rs34903665 | 0.6885 | 1/1 | 0/1 | 0/1 | 0/1 | 1/1 | 0/0 | 0/1 | 1/1 | 0/0 | 0/1 | 0/1 | 0/0 | |
| 38945169 | A | G | FAM114A1 | exonic | rs1060582 | 0.1114 | 0/1 | 0/0 | 0/0 | 0/0 | 0/0 | 0/0 | 0/0 | 0/0 | 0/0 | 0/0 | 0/0 | 0/0 | |
| 38972691 | T | C | TMEM156 | exonic | rs140693293 | 0.0038 | 0/0 | 0/0 | 0/0 | 0/0 | 0/0 | 0/0 | 0/0 | 0/0 | 0/0 | 0/1 | 0/0 | 0/0 | |
| 38972793 | A | G | TMEM156 | intronic | rs3733268 | 0.0197 | 0/0 | 0/1 | 0/0 | 0/0 | 0/0 | 0/0 | 0/0 | 0/0 | 0/0 | 0/0 | 0/0 | 0/0 | |
| 38990334 | C | T | TMEM156 | intronic | rs3821986 | 0.4426 | 0/1 | 1/1 | 1/1 | 0/1 | 1/1 | 0/0 | 1/1 | 0/0 | 0/1 | 1/1 | 1/1 | 0/0 | |
| 38995310 | A | G | TMEM156 | intronic | rs11721954 | 0.3605 | 0/0 | 0/0 | 0/1 | 0/0 | 0/0 | 0/0 | 0/1 | 0/0 | 0/1 | 1/1 | 1/1 | 0/0 | |
| 38995374 | T | C | TMEM156 | exonic | rs10212770 | 0.88 | 0/1 | 1/1 | 1/1 | 1/1 | 1/1 | 1/1 | 1/1 | 1/1 | 1/1 | 1/1 | 1/1 | 0/0 | **C** |
| 39000305 | A | G | TMEM156 | exonic | rs11542133 | 0.23 | 0/0 | 0/0 | 0/0 | 0/0 | 0/0 | 0/0 | 0/0 | 0/0 | 0/1 | 0/1 | 0/0 | 0/0 | **A** |
| 39064126 | G | C | KLHL5 | UTR5 | rs2711942 | 0.74 | 0/1 | 1/1 | 1/1 | 1/1 | 1/1 | 0/0 | 0/0 | 1/1 | 1/1 | 1/1 | 1/1 | 1/1 | **C** |
| 39064162 | A | C | KLHL5 | exonic | rs2711941 | 0.74 | 0/1 | 1/1 | 1/1 | 1/1 | 1/1 | 0/0 | 0/0 | 1/1 | 1/1 | 1/1 | 1/1 | 1/1 | **C** |
| 39077567 | G | GT | KLHL5 | intronic | rs201873328 | 0.02 | 0/0 | 0/0 | 0/0 | 0/0 | 0/0 | 0/0 | 0/0 | 0/0 | 0/1 | 0/0 | 0/0 | 0/0 | **G** |
| 39114929 | G | A | KLHL5 | intronic | rs3796510 | 0.53 | 0/0 | 0/0 | 0/0 | 0/0 | 0/0 | 0/0 | 0/1 | 0/0 | 0/0 | 0/0 | 0/0 | 1/1 | **G** |
| 39116775 | A | T | KLHL5 | intronic | rs10026775 | 0.53 | 0/0 | 0/0 | 0/0 | 0/0 | 0/0 | 0/0 | 0/1 | 0/0 | 0/0 | 0/0 | 0/0 | 1/1 | **A** |
| 39116911 | T | C | KLHL5 | exonic | rs3733276 | 0.53 | 0/0 | 0/0 | 0/0 | 0/0 | 0/0 | 0/0 | 0/1 | 0/0 | 0/0 | 0/0 | 0/0 | 1/1 | **T** |
| 39117103 | G | A | KLHL5 | intronic | rs3733277 | 0.53 | 0/0 | 0/0 | 0/0 | 0/0 | 0/0 | 0/0 | 0/1 | 0/0 | 0/0 | 0/0 | 0/0 | 1/1 | **G** |
| 39205365 | C | T | WDR19 | intronic | rs1451817 | 0.97 | 1/1 | 1/1 | 1/1 | 1/1 | 1/1 | 1/1 | 1/1 | 1/1 | 1/1 | 1/1 | 1/1 | 1/1 | **T** |
| 39216221 | C | T | WDR19 | exonic | rs2167494 | 0.29 | 0/1 | 0/0 | 0/0 | 0/0 | 0/0 | 0/0 | 0/1 | 0/0 | 0/0 | 0/0 | 0/0 | 0/0 | **C** |
| 39216240 | G | A | WDR19 | exonic | rs75964850 | 0.04 | 0/0 | 0/0 | 0/0 | 0/0 | 0/0 | 0/0 | 0/1 | 0/0 | 0/0 | 0/0 | 0/0 | 0/0 | **G** |
| 39217779 | C | T | WDR19 | exonic | rs199765304 | 0 | 0/0 | 0/0 | 0/0 | 0/0 | 0/0 | 0/0 | 0/1 | 0/0 | 0/0 | 0/0 | 0/0 | 0/0 | **A** |
| 39229771 | G | A | WDR19 | intronic | rs11730558 | 0.31 | 0/1 | 0/0 | 0/0 | 0/0 | 0/0 | 0/0 | 0/1 | 0/0 | 0/0 | 0/0 | 0/0 | 0/0 | **G** |
| 39242111 | C | A | WDR19 | intronic | rs9998591 | 0.64 | 0/1 | 1/1 | 1/1 | 1/1 | 1/1 | 1/1 | 0/1 | 1/1 | 1/1 | 1/1 | 1/1 | 1/1 | **A** |
| 39254828 | A | C | WDR19 | exonic | rs187546086 | 0 | 0/1 | 0/0 | 0/0 | 0/0 | 0/0 | 0/0 | 0/0 | 0/0 | 0/0 | 0/0 | 0/0 | 0/0 | **A** |
| 39255496 | A | G | WDR19 | intronic | rs33997556 | 0.08 | 0/1 | 1/1 | 1/1 | 1/1 | 1/1 | 1/1 | 0/1 | 1/1 | 1/1 | 1/1 | 1/1 | 0/0 | **G** |
| 39271541 | A | G | WDR19 | intronic | rs3733280 | 0.31 | 0/1 | 0/0 | 0/0 | 0/0 | 0/0 | 0/0 | 0/1 | 0/0 | 0/0 | 0/0 | 1/1 | 0/0 | **A** |
| 39276623 | A | AG | WDR19 | intronic | rs11096989 | 0.5 | 1/1 | 1/1 | 1/1 | 1/1 | 1/1 | 1/1 | 1/1 | 1/1 | 1/1 | 1/1 | 1/1 | 0/0 | **AG** |
| 39279724 | T | C | WDR19 | intronic | rs12648082 | 0.5 | 1/1 | 1/1 | 1/1 | 1/1 | 1/1 | 1/1 | 1/1 | 1/1 | 1/1 | 1/1 | 1/1 | 0/0 | **C** |
| 39279907 | A | G | WDR19 | intronic | rs2276888 | 0.46 | 0/0 | 0/0 | 0/0 | 0/0 | 0/0 | 0/0 | 0/0 | 0/0 | 0/0 | 0/0 | 0/0 | 1/1 | **A** |
| 39297211 | C | T | RFC1 | intronic | rs41547922 | 0 | 0/0 | 0/0 | 0/1 | 0/1 | 0/0 | 0/0 | 0/0 | 0/0 | 0/0 | 0/0 | 0/0 | 0/0 | **C** |
| 39301605 | G | C | RFC1 | intronic | rs2066788 | 0.51 | 1/1 | 1/1 | 1/1 | 1/1 | 1/1 | 1/1 | 1/1 | 1/1 | 1/1 | 1/1 | 1/1 | 0/0 | **C** |
| 39302029 | T | C | RFC1 | exonic | rs2066786 | 0.45 | 0/0 | 0/0 | 0/0 | 0/0 | 0/0 | 0/0 | 0/0 | 0/0 | 0/0 | 0/0 | 0/0 | 1/1 | **T** |
| 39303925 | A | G | RFC1 | exonic | rs2066782 | 0.1 | 1/1 | 1/1 | 1/1 | 1/1 | 1/1 | 1/1 | 0/1 | 1/1 | 1/1 | 1/1 | 0/0 | 0/0 | **G** |
| 39329102 | G | A | RFC1 | intronic | rs11096992 | 0.47 | 1/1 | 1/1 | 1/1 | 1/1 | 1/1 | 1/1 | 1/1 | 1/1 | 1/1 | 1/1 | 1/1 | 1/1 | **A** |
| 39353122 | C | T | RFC1 | intronic | rs4975007 | 0.98 | 1/1 | 1/1 | 1/1 | 1/1 | 1/1 | 1/1 | 1/1 | 1/1 | 1/1 | 1/1 | 1/1 | 1/1 | **T** |
| 39353137 | C | T | RFC1 | intronic | rs374311239 | 0 | 0/0 | 0/1 | 0/0 | 0/0 | 0/0 | 0/0 | 0/0 | 0/0 | 0/0 | 0/0 | 0/0 | 0/0 | **C** |
| 39439264 | T | C | KLB | intronic | rs4975015 | 0.1707 | 0/1 | 0/0 | 0/1 | 0/0 | 0/0 | 0/0 | 1/1 | 1/1 | 1/1 | 0/0 | 0/0 | 1/1 | **C** |
| 39448529 | G | A | KLB | exonic | rs17618244 | 0.1528 | 0/0 | 0/1 | 0/0 | 0/0 | 1/1 | 0/0 | 0/0 | 0/0 | 0/0 | 0/1 | 1/1 | 0/0 | **G** |
| 39448542 | C | G | KLB | exonic | rs7685429 | 0.7326 | 1/1 | 1/1 | 0/1 | 1/1 | 1/1 | 1/1 | 0/0 | 1/1 | 1/1 | 0/1 | 1/1 | 0/0 | **G** |
| 39448586 | C | T | KLB | exonic | rs35372803 | 0.0375 | 0/0 | 0/1 | 0/0 | 0/0 | 0/0 | 0/0 | 0/0 | 0/0 | 0/0 | 0/0 | 0/0 | 0/0 | **C** |
| 39450229 | C | A | KLB | exonic | rs4975017 | 0.2881 | 0/1 | 0/0 | 0/0 | 0/0 | 0/0 | 0/0 | 0/0 | 0/0 | 1/1 | 0/0 | 0/0 | 0/0 | |
| 39450403 | TGA | T | KLB | UTR3 | rs112327399 | 0.0603 | 0/0 | 0/0 | 0/0 | 0/0 | 0/0 | 0/0 | 0/0 | 1/1 | 0/0 | 0/0 | 0/0 | 0/0 | |
| 39457857 | T | C | RPL9 | intronic | rs1015450 | 0.1489 | 0/1 | 0/1 | 0/0 | 0/1 | 0/0 | 0/0 | 0/0 | 0/0 | 0/0 | 0/0 | 0/0 | 0/0 | |
| 39457870 | T | TG | RPL9 | intronic | rs3216720 | 0.6465 | 1/1 | 1/1 | 0/1 | 1/1 | 1/1 | 1/1 | 0/0 | 0/0 | 1/1 | 0/1 | 1/1 | 0/0 | |
| 39457954 | G | C | RPL9 | intronic | rs117602443 | 6.00E-04 | 0/0 | 0/0 | 0/0 | 0/0 | 0/0 | 0/0 | 0/0 | 0/0 | 0/0 | 0/1 | 0/0 | 0/0 | |
| 39459724 | T | C | RPL9 | intronic | rs2687958 | 0.6479 | 1/1 | 1/1 | 0/1 | 1/1 | 1/1 | 1/1 | 0/0 | 0/0 | 1/1 | 0/1 | 1/1 | 0/0 | |
| 39460328 | G | A | RPL9 | intronic | rs73137448 | 0.0563 | 0/0 | 0/0 | 0/0 | 0/0 | 0/0 | 0/0 | 0/0 | 1/1 | 0/0 | 0/0 | 0/0 | 0/0 | |
| 39460352 | C | G | RPL9 | intronic | rs11550000 | 0.2497 | 0/1 | 0/1 | 0/0 | 0/1 | 0/0 | 0/0 | 0/0 | 1/1 | 0/0 | 0/0 | 0/0 | 0/0 | |
| 39460413 | C | A | RPL9 | intronic | rs2276889 | 0.0234 | 0/0 | 0/0 | 0/1 | 0/1 | 0/0 | 1/1 | 0/0 | 0/0 | 0/0 | 0/0 | 0/0 | 0/0 | |
| 39460490 | C | G | RPL9 | intronic | rs2276890 | 0.2844 | 0/1 | 0/0 | 0/0 | 0/0 | 0/0 | 0/0 | 0/0 | 0/0 | 1/1 | 0/0 | 0/0 | 0/0 | |
| 39460539 | G | A | RPL9 | UTR5 | rs116433592 | 0.005 | 0/1 | 0/0 | 0/0 | 0/0 | 0/0 | 0/0 | 0/0 | 0/0 | 0/0 | 0/0 | 0/0 | 0/0 | |
| 39460572 | G | A | LIAS\x3bRPL9 | upstream | rs3821989 | 0.006 | 0/0 | 0/0 | 0/0 | 0/0 | 0/0 | 0/0 | 0/0 | 0/0 | 0/0 | 0/1 | 0/0 | 0/0 | |
| 39460797 | G | A | LIAS | intronic | rs2687959 | 0.1826 | 0/0 | 0/1 | 0/1 | 0/1 | 1/1 | 1/1 | 0/0 | 0/0 | 0/0 | 0/1 | 1/1 | 0/0 | |
| 39466639 | A | G | LIAS | intronic | rs76534325 | 0.0378 | 0/0 | 0/0 | 0/0 | 0/0 | 0/0 | 0/0 | 0/0 | 1/1 | 0/0 | 0/0 | 0/0 | 0/0 | |
| 39469279 | A | T | LIAS | intronic | rs79572060 | 0.0233 | 0/0 | 0/0 | 0/1 | 0/1 | 0/0 | 1/1 | 0/0 | 0/0 | 0/0 | 0/0 | 0/0 | 0/0 | |
| 39472786 | C | T | LIAS | intronic | rs79140300 | 0.0308 | 0/0 | 0/0 | 0/1 | 0/1 | 0/0 | 1/1 | 0/0 | 0/0 | 0/0 | 0/0 | 0/0 | 0/0 | |
| 39472796 | T | A | LIAS | intronic | rs73137454 | 0.0747 | 0/0 | 0/0 | 0/0 | 0/0 | 0/0 | 0/0 | 0/0 | 1/1 | 0/0 | 0/0 | 0/0 | 0/0 | |
| 39472842 | T | A | LIAS | intronic | rs2125314 | 0.3142 | 0/1 | 0/0 | 0/0 | 0/0 | 0/0 | 0/0 | 0/0 | 0/0 | 1/1 | 0/0 | 0/0 | 0/0 | |
| 39474868 | G | A | LIAS | intronic | rs749915 | 0.236 | 0/1 | 0/1 | 0/0 | 0/1 | 0/0 | 0/0 | 0/0 | 1/1 | 0/0 | 0/0 | 0/0 | 0/0 | |
| 39482384 | A | G | LOC401127 | ncRNA exonic | rs10009841 | 0.0747 | 0/0 | 0/0 | 0/0 | 0/0 | 0/0 | 0/0 | 0/0 | 1/1 | 0/0 | 0/0 | 0/0 | 0/0 | |
| 39482440 | G | A | LOC401127 | ncRNA exonic | rs2276891 | 0.0232 | 0/0 | 0/0 | 0/1 | 0/1 | 0/0 | 1/1 | 0/0 | 0/0 | 0/0 | 0/0 | 0/0 | 0/0 | |
| 39482584 | T | C | LOC401127 | ncRNA exonic | rs17438520 | 0.0259 | 0/1 | 0/0 | 0/0 | 0/0 | 0/0 | 0/0 | 0/0 | 0/0 | 1/1 | 0/0 | 0/0 | 0/0 | |
| 39482627 | T | C | LOC401127 | ncRNA exonic | rs2276893 | 0.0232 | 0/0 | 0/0 | 0/1 | 0/1 | 0/0 | 1/1 | 0/0 | 0/0 | 0/0 | 0/0 | 0/0 | 0/0 | |
| 39483115 | A | G | LOC401127 | ncRNA exonic | rs2687980 | 0.7109 | 1/1 | 1/1 | 0/1 | 1/1 | 1/1 | 1/1 | 0/0 | 1/1 | 1/1 | 0/1 | 1/1 | 0/0 | |
| 39506144 | C | T | UGDH | intronic | rs1379819 | 0.1465 | 0/0 | 0/0 | 0/0 | 0/0 | 0/0 | 0/0 | 0/0 | 0/0 | 0/0 | 1/1 | 0/0 | 0/0 | |
| 39506997 | GA | G | UGDH | intronic | rs33942301 | 0.248 | 0/1 | 0/0 | 0/1 | 0/1 | 0/0 | 1/1 | 0/0 | 0/0 | 1/1 | 1/1 | 0/0 | 0/0 | |
| 39507083 | G | T | UGDH | intronic | rs4975019 | 0.3746 | 0/1 | 0/0 | 0/1 | 1/1 | 0/0 | 1/1 | 0/0 | 0/0 | 1/1 | 1/1 | 0/0 | 0/0 | |
| 39510175 | G | A | UGDH | intronic | rs6815780 | 0.2211 | 0/1 | 0/0 | 0/1 | 0/1 | 0/0 | 1/1 | 0/0 | 0/0 | 1/1 | 1/1 | 0/0 | 0/0 | |
| 39510264 | A | G | UGDH | exonic | rs1129052 | 0.2203 | 0/1 | 0/0 | 0/1 | 0/1 | 0/0 | 1/1 | 0/0 | 0/0 | 1/1 | 1/1 | 0/0 | 0/0 | |
| 39511359 | T | C | UGDH | intronic | rs6531722 | 0.2211 | 0/1 | 0/0 | 0/1 | 0/1 | 0/0 | 1/1 | 0/0 | 0/0 | 1/1 | 1/1 | 0/0 | 0/0 | |
| 39512347 | T | C | UGDH | exonic | rs11544855 | 0.1145 | 0/0 | 0/0 | 0/1 | 0/0 | 0/0 | 0/0 | 1/1 | 0/0 | 0/0 | 0/0 | 0/0 | 1/1 | |
| 39515736 | A | G | UGDH | exonic | rs10019532 | 0.213 | 0/1 | 0/0 | 0/1 | 0/1 | 0/0 | 1/1 | 0/0 | 0/0 | 1/1 | 1/1 | 0/0 | 0/0 | |
| 39515806 | TAAAAAGA | T | UGDH | intronic | rs151116497 | 0.1451 | 0/0 | 0/0 | 0/0 | 0/0 | 0/0 | 0/0 | 0/0 | 0/0 | 0/0 | 1/1 | 0/0 | 0/0 | |
| 39528752 | G | A | UGDH | intronic | rs4975020 | 0.7065 | 1/1 | 0/1 | 1/1 | 1/1 | 0/0 | 1/1 | 1/1 | 0/0 | 1/1 | 1/1 | 1/1 | 1/1 | |
| 39606900 | T | C | SMIM14 | intronic | rs147611359 | 0.0076 | 0/0 | 0/0 | 0/0 | 0/0 | 0/0 | 0/0 | 0/1 | 0/0 | 0/0 | 0/0 | 0/0 | 0/0 | |
| 39757480 | A | AT | UBE2K | intronic | rs11389468 | 0.8191 | 1/1 | 1/1 | 0/1 | 1/1 | 0/0 | 1/1 | 0/1 | 1/1 | 0/1 | 1/1 | 1/1 | 1/1 | |
| 39757525 | C | T | UBE2K | intronic | rs138858087 | 0.0072 | 0/0 | 0/0 | 0/0 | 0/0 | 0/0 | 0/0 | 0/1 | 0/0 | 0/0 | 0/0 | 0/0 | 0/0 | |
| 39871117 | G | GA | PDS5A | intronic | rs34039605 | 0.3225 | 0/1 | 1/1 | 0/0 | 0/1 | 0/0 | 0/0 | 0/0 | 1/1 | 0/1 | 1/1 | 1/1 | 0/0 | |
| 39900041 | T | C | PDS5A | exonic | rs193075238 | 3.24E-05 | 0/0 | 0/0 | 0/0 | 0/0 | 0/0 | 0/0 | 0/0 | 0/0 | 0/0 | 0/0 | 1/1 | 0/0 | |
| 39903924 | G | A | PDS5A | intronic | rs77174235 | 0.0728 | 0/1 | 0/1 | 0/0 | 0/1 | 0/0 | 0/0 | 0/0 | 1/1 | 0/1 | 0/0 | 0/0 | 0/0 | |

### Table S4. CANVAS core haplotype from WES data (core haplotype shown in dark green)
